# Immunomodulatory effect of mycobacterial outer membrane vesicles coated nanoparticles

**DOI:** 10.1101/2022.06.07.495204

**Authors:** Edna George, Avijit Goswami, Tejan Lodhiya, Priyanka Padwal, Shalini Iyer, Iti Gauttam, Lakshay Sethi, Sharumathi Jeyasankar, Pallavi Raj Sharma, Ameya Atul Dravid, Raju Mukherjee, Rachit Agarwal

**Affiliations:** Centre for BioSystems Science and Engineering, Indian Institute of Science, Bangalore, India; Indian Institute of Science Education and Research, Tirupati, India

**Author notes:** Equal contribution (Edna George, Avijit Goswami).

## Abstract

Tuberculosis (TB) is one of the most widely prevalent infectious diseases that cause significant mortality. Bacillus Calmette-Guérin (BCG), the current TB vaccine used in clinics, shows variable efficacy and has safety concerns for immunocompromised patients. There is a need to develop new and more effective TB vaccines. Outer membrane vesicles (OMVs) are vesicles released by Mycobacteria that contain several lipids and membrane proteins and act as a good source of antigens to prime immune response. However, the use of OMVs as vaccines has been hampered by their heterogeneous size and low stability. Here we report that mycobacterial OMVs can be stabilized by coating over uniform-sized 50 nm gold nanoparticles. The OMV-coated gold nanoparticles (OMV-AuNP) show enhanced uptake and activation of macrophages and dendritic cells. Proteinase K and TLR inhibitor studies demonstrated that the enhanced activation was attributed to proteins present on OMVs and was mediated primarily by TLR2 and TLR4. Mass spectrometry analysis revealed several potential membrane proteins that were common in both free OMVs and OMV-AuNP. Such strategies may open up new avenues and the utilization of novel antigens for developing TB vaccines.

## Introduction

Tuberculosis (TB) is a significant health concern due to its pathogenicity and mortality. As a pathogen, *Mycobacterium tuberculosis* (Mtb) has infected millions of people worldwide and has contributed to one of the leading causes of global deaths [1]. Around 1.5 million deaths due to TB were reported in 2020 (Global Tuberculosis Report 2021, World Health Organization). Hence, there is an urgent need to establish an effective preventive treatment to curtail the spread of TB. Vaccines are an integral part of public health to protect the population and control the spread of various communicable diseases. Bacillus Calmette-Guérin (BCG) is the only clinically available vaccine against TB with protection limited to the pediatric group [2]. Additionally, BCG-associated complications due to immunodeficiency or HIV infection (ones who need the vaccine the most) restricts its use [3-5]. This highlights the urgent need for new vaccine strategies that ensure protection throughout adulthood. Vaccine development has been an area of active research to substitute the current BCG vaccine with potent antigens and efficient delivery systems [6] or by modifying the genetic background of the causative pathogen [7]. **Error! Bookmark not defined**.For example, MVA85A (a subunit vaccine) and MTBVAC (a live attenuated vaccine) have advanced to clinical trials [8]. However, concerns around using these vaccines in immunocompromised patients and pre-existing immunity to viral vectors still exist [3].

The efficacy of a vaccine rests in identifying suitable antigens to trigger the immune system. Several studies have been carried out to explore the possibility of using immunogenic proteins like Antigen 85, ESAT6, and CFP10 to develop a potential vaccine for TB [9-11]. While promising, vaccines using these antigens did not prevent infection completely [9, 12]. Hence, there is a need to develop new approaches and antigens for TB vaccines. Bacterial outer membrane vesicles (OMVs) composed of proteins, lipids, and glycans are nanosized structures (20-250 nm) naturally released by bacteria [13]. These vesicles serve several functional roles that include secretion of virulence factors [14], delivery or transport of bacterial products [15, 16], and signaling [17]. The surface proteins on OMVs also contribute to the pathogenicity of the microbes and can be stimulating for humoral and cell-mediated immune responses in the host [18, 19]. Lipoproteins and glycolipids on the OMVs act as pathogen-associated molecular patterns (PAMPs) that interact with host cells. These are recognized by pattern recognition receptors (PRRs) on the host cells, eliciting a pro-inflammatory response in the host, mediated primarily by Toll-like receptors (TLRs) [20]. In addition, several membrane proteins and lipid antigens are also displayed on OMV surfaces that are known to be important for the induction of protective immunity [21-24].

Several studies have been carried out to understand the host-OMV interactions to exploit the full potential of OMVs. OMV vaccine against *Neisseria meningitidis* is clinically approved and has shown efficacy against serogroup B meningococcal disease [25]. Designer OMVs have also been developed where antigens are engineered on OMV surfaces to induce a robust immune response [26, 27]. Earlier studies have shown the protective capacity of outer membrane vesicles (OMV) isolated from *Mycobacterium smegmatis (M. smegmatis)* [28, 29] or *Mycobacterium bovis* [30] against TB. Alum adjuvanted OMVs from *M. smegmatis* and OMVs from *Mycobacterium bovis* were shown to elicit cross-reactive humoral and cellular responses against *M. tuberculosis* antigens [29, 31] and reduced bacterial load in mice after *M. tuberculosis* challenge [28, 30]. Despite promising results, some disadvantages such as compromised vesicle integrity and stability and heterogeneous size resulting in variable pharmacokinetics are associated with free OMVs [32-34]. Combining OMVs with synthetic nanoparticles can overcome some of the disadvantages of free OMVs. The use of nanoparticles allows control over various physicochemical parameters such as size, shape, stability, etc. Wu *et al*. showed that coating OMVs over 100 nm bovine serum albumin particles induced higher humoral and cellular immunity against carbapenem-resistant *Klebsiella pneumoniae* than equivalent free OMVs [33]. Gao *et al*. coated OMVs isolated from *Escherichia coli* over 30 nm gold nanoparticles and showed more potent specific antibody and T-cell response compared to free OMVs [34]. However, studies involving nanoparticles coated with OMVs from Mycobacteria and their host-microbe interaction remain elusive. In this study, we aim to coat mycobacterial OMVs on gold nanoparticles to modulate the immune response and understand the interaction of OMVs with innate immune cells. We found that OMVs can be coated over gold nanoparticles and form a stable formulation of defined size. Interestingly, OMV coated gold nanoparticles showed several-fold enhanced immune-modulatory effects than free OMVs. This approach can open new avenues to explore mycobacterial OMVs for vaccines against TB.

## Materials and Methods

### Synthesis and characterization of gold nanoparticles (AuNP)

Gold nanoparticles (AuNP) were synthesized by a one-pot seeded growth method with tris-base [35]. Initially, gold seeds were synthesized by adding 0.5 mL of 25 mM HAuCl_4_ (Sigma) to 48 mL of boiling water and continued heating for another 10 min under stirring. Next, 1.5 mL of 1% sodium citrate (Sigma) was added to the boiling HAuCl_4_ solution. The color of the solution changed from yellow to bluish-grey to ruby red. The reaction was continued for 30 min to obtain monodispersed gold seeds. For synthesizing AuNP using the seeded growth method, 2 mL of 0.1 M tris-base was added to 45.5 mL of boiling water and left for stirring for another 5 min. Two mL of gold seeds were added to 0.5 mL of 25 mM HAuCl_4_ solution. The reaction was continued for 30 min under heated stirring.

To characterize the synthesized nanoparticles, the reaction was cooled, and samples were taken for recording the absorption spectra (Tecan Infinite Pro). The sizes of the seeds were characterized using dynamic light scattering (DLS, Malvern Zetasizer). The particle size of the resulting AuNP was initially characterized using DLS. For transmission electron microscopy (TEM), samples were deposited on a copper grid and rinsed after 5 min with distilled water before being visualized using 300kV Tecnai™ F30 TEM/STEM microscope.

### Bacterial culture and isolation of outer membrane vesicles (OMVs)

OMVs were isolated from *M. smegmatis* according to the standard protocols [13]. Briefly, *M. smegmatis* cell suspension at the late log phase was centrifuged at 4000 g to remove cells and large cellular debris. The supernatant was collected, sheared using a 22-gauge needle, filtered using 0.45 µm pore-sized polyethersulfone syringe filters, followed by 0.22 µm pore-sized cellulose acetate syringe filters, and concentrated using 100 kDa concentrators to concentrate the OMVs. Later, OMVs were purified and further concentrated by ultracentrifugation (Beckman L8-60M, Ti-50 rotor) at 150,000 g for two hours at 4°C. OMVs pellet was washed twice, resuspended in 150 µL water, and stored at -80°C until further use. The protein content was determined by bicinchoninic acid (BCA) assay, and the sizes of OMVs were characterized using DLS measurements.

### Synthesis and characterization of outer membrane vesicles coated nanoparticles (OMV-AuNP)

To coat gold nanoparticles with OMVs, 50 μg/mL of AuNP were mixed with the 150 µg/mL OMVs isolated from the non-pathogenic *M. smegmatis* and then co-extruded nine times through a 100 nm pore size polycarbonate porous membrane using a mini extruder (Avanti Polar Lipids). Excess OMVs and soluble compounds were removed from the supernatant after centrifugation at 15000 g for 10 minutes at 4°C. The final concentration of the nanoparticles was measured by plotting the absorbance at 530 nm (Tecan Infinite Pro) against a standard curve of AuNP. Transmission electron microscopy (TEM) was used to observe the morphology of these OMV coated nanoparticles (OMV-AuNP). These particles were deposited onto a glow discharged copper grid and rinsed with distilled water, followed by 1% uranyl acetate staining. Subsequently, the grid was dried and visualized using TEM. Long-term stability was examined by monitoring the size in water using Zetasizer. For quantifying protein content on OMV-AuNP using SDS-PAGE, extruded OMV-AuNP were run on a 10% polyacrylamide gel and visualized by silver staining (Pierce Silver staining Kit, #24612) using manufacturer’s protocol. Protein gel bands were quantified using ImageJ software. Relative band intensities were used to plot a standard curve corresponding to respective known free OMV protein concentration, which was then used to quantify the absolute protein content after extrusion. Additionally, OMV coating on AuNP was determined by quantifying the protein content in OMV using a CBQCA protein quantitation kit (ThermoFisher Scientific). For stability studies of OMV coated AuNP, nanoparticles and their respective controls were stored at 4°C. The sizes of these samples were observed using DLS and transmittance was calculated using absorbance measurements at 560 nm on days 0, 1, 3, 5, 7, 14, and 22 using DLS measurements.

### Cell culture

Human monocytic leukemia cell line, THP-1 cells, were cultured in 25 cm^2^ flasks in RPMI media (Gibco) supplemented with 10% fetal bovine serum (Sigma) and 1% penicillin/streptomycin (Gibco). These cells were maintained in a CO_2_ incubator (Thermo Scientific) with 95% humidified air and 5% CO_2_ at 37°C. The cells were passaged every third day or when the culture flasks were 70-80% confluent.

### Mice

C57BL/6 mice were used for experiments with approval from the Institutional Animal Ethics Committee (CAF/Ethics/733/2020). These mice were housed and maintained under standard laboratory conditions in the Central Animal Facility at the Indian Institute of Science, Bangalore. Both male and female mice were used in this study.

### Isolation of macrophages and dendritic cells from mouse bone marrow

Bone marrow cells were flushed from femurs and tibias of mice under aseptic conditions. The cells were washed once with basal media, treated with RBC lysis buffer (0.8% NH_4_Cl in 10 mM Tris buffer, pH 7.5), and washed again with basal media. The cell numbers were adjusted to the requirement of the experiment, cultured in RPMI media (Gibco) supplemented with 10% fetal bovine serum (Sigma), growth factor, and 1% penicillin/streptomycin (Gibco), and maintained in a CO_2_ incubator at 37°C with 95% humidified air and 5% CO_2._ Depending on the requirement of bone marrow-derived macrophages (BMDM) or bone marrow-derived dendritic cells (BMDC), the growth factors M-CSF or GM-CSF (20 ng/mL, BioLegend) was added to the above media, respectively.

### Cell uptake studies of OMV-AuNP

THP-1 monocytes were plated on coverslips at a seeding density of 30000 cells/well in a 24-well plate and differentiated with 20 ng/mL phorbol myristate acetate (Sigma). After 18 h, the cells were washed, replenished with fresh media, and rested for 48 h. For cell uptake studies, OMVs were incubated with DiI (Thermo) at a 1:1000 ratio for 30 mins at room temperature to make them fluorescent. These fluorescent OMVs were extruded with AuNPs and washed twice to remove excess stain and unbound OMVs. Finally, the cells were washed twice and incubated with OMV-AuNP at a final AuNP concentration of 20 µg/mL. Cells treated with free OMVs, unextruded AuNP with free OMVs, and untreated cells served as controls. After incubating the cells for 24 h, the cells were stained for actin (Thermo Molecular Probes) and imaged using an InCell microscope (GE InCell Analyzer 6000). A minimum of 25 cells per well were analyzed for particle uptake by quantifying the mean fluorescence from DiI using ImageJ (NIH).

### Proteinase K assay and TLR2/4 inhibition studies

20 µg of OMV-AuNP was treated with or without 20 µg of proteinase K at 37°C for 20 mins. After incubation, excess proteinase K was removed by centrifugation at 15000 g for 10 mins, and the OMV-AuNP pellet was washed twice with water. 20 µg/mL of OMV-AuNP treated or untreated with proteinase K was added to THP-1 cells and incubated for 24 h in a CO_2_ incubator at 37°C with 95% humidified air and 5% CO_2_. Cells were processed for RNA isolation to quantify cytokine mRNA levels using RT-PCR.

### Cytokine mRNA studies

For mRNA studies, RNA was isolated using Tri Reagent (Sigma). 600 ng of RNA was used for cDNA conversion using the manufacturer’s protocol below (High Capacity cDNA reverse transcriptase kit).

### 15 µL Reaction

RNA to cDNA conversion

15 µL Reaction: RNA to cDNA conversion
|  |  |
| --- | --- |
| 10X RT Buffer | 1.5 µL |
| Random primers | 1.5 µL |
| dNTPs Mix (100 mM) | 0.8 µL |
| Reverse Transcriptase (50 U/µL) | 1 µL |
| RNA | 600ng |
| Milli Q | Makeup volume 15 µL |

### Reaction temperature cycle

Reaction temperature cycle:
|  |  |
| --- | --- |
| 25° C | 10 mins |
| 37 ° C | 120 mins |
| 85° C | 5 mins |
| 4° C | Hold |

cDNA was diluted four times with ultrapure H_2_O, and PCR amplification was done using gene-specific primers with iTaq SYBR green (BioRad) in CFX 96 real-time PCR machine. List of primers used is present in **Supplementary Table 1**.

**Table 1:** *M. smegmatis mc*^*2*^*155* proteins commonly identified both in free-OMVs and OMV-AuNPs. PSM and protein FDR < 0.01. Proteins were identified with at least 1 unique peptide.

| Protein IDs | H37Rv Ortholog | Functional category | Function |
| --- | --- | --- | --- |
| MSMEG_3618 | Rv1860 | Cell wall and cell processes | Fibronectin attachment protein<br>immunogenic protein MPT32 |
| MSMEG_3637 | Rv1842c | Cell wall and cell processes | Conserved hypothetical membrane protein |
| MSMEG_6120 | #N/A | Cell wall and cell processes | ABC transporter |
| MSMEG_6195 | Rv3680 | Cell wall and cell processes | Probable anion transporter<br>ATPase |
| MSMEG_0306 | Rv3566c | Intermediary metabolism and respiration | Arylamine N-acetyltransferase<br>Nat (arylamine acetylase) |
| MSMEG_0485 | #N/A | Intermediary metabolism and respiration | Amidase |
| MSMEG_0925 | #N/A | Intermediary metabolism and respiration | SAM-dependent methyltransferase |
| MSMEG_2681 | Rv0840c | Intermediary metabolism and respiration | Probable proline iminopeptidase Pip (prolyl aminopeptidase) (pap) |
| MSMEG_3306 | #N/A | Intermediary metabolism and respiration | Zinc-binding alcohol dehydrogenase |
| MSMEG_4862 | #N/A | Intermediary metabolism and respiration | 3-alpha-hydroxysteroid dehydrogenase |
| MSMEG_1554 | #N/A | Conserved hypotheticals | Ethanolamine ammonia-lyase |
| MSMEG_2226 | #N/A | Conserved hypotheticals | Unknown |
| MSMEG_3374 | #N/A | Conserved hypotheticals | Unknown |
| MSMEG_6675 | #N/A | Conserved hypotheticals | Unknown |
| MSMEG_1140 | Rv0564c | Lipid metabolism | Probable glycerol-3-phosphate dehydrogenase |
| MSMEG_3139 | Rv1472 | Lipid metabolism | Possible enoyl-CoA hydratase<br>EchA12 (enoyl hydratase) |
| MSMEG_1365 | Rv0652 | Information pathways | 50S ribosomal protein L7/L12<br>RplL (SA1) |
| MSMEG_2489 | #N/A | Regulatory proteins | Transcriptional regulator,<br>GntR |

### Reaction temperature cycle

| Temp | Time | Cycle |
| --- | --- | --- |
| 95° C | 10 mins |  |
| 95 ° C | 20 secs | 40 cycles |
| 55° C | 25 secs |  |
| 72 °C | 30 secs |  |

### Evaluation of antigenic properties of OMV-AuNP

THP-1 cells plated at a seeding density of 35000 cells/well in a 96-well plate were activated with 20 ng/mL phorbol 12-myristate 13-acetate (Sigma) and incubated. After 18 h, the cells were washed and replenished with fresh media and rested for 48 h. For studies on BMDM, cells were seeded at 5 × 10^5^ cells/ well in a 48-well plate. The adherent cells were washed twice and incubated with free OMVs, uncoated AuNP, and OMV-AuNP at a final AuNP concentration of 20 µg/mL. Untreated cells and cells treated with LPS (1 µg/mL) were used as negative and positive controls. The media collected from each well after 24 h of incubation was used directly or stored at -80°C until further use. For quantifying the cytokines released from THP-1 cells, hIL-8 and hIL-6 were estimated using their respective ELISA kits (Peprotech) according to the manufacturer’s instructions. Similarly, for profiling the cytokines released from BMDM, levels of mIL-6 and mTNF-α were estimated using their respective ELISA kits (ThermoFisher Scientific).

### Proteomics sample preparation

#### Reagents

All reagents were procured from Merk. Trypsin (sequencing grade modified, porcine) was procured from Promega. Optima LC/MS grade solvents, water, acetonitrile, and formic acid were procured from Fisher chemicals.

#### Protein digestion and sample preparation

The precipitated mixture of protein was reconstituted in 6 M urea at 35°C with constant shaking. Five µg of protein were subsequently reduced with 1,4-Dithiothreitol (DTT) and alkylated with iodoacetamide with a final concentration of 10 mM and 37 mM, respectively. The reactions were quenched with 30mM DTT. The mixture of linearized proteins was digested with trypsin (1:5 enzyme/proteins) for 16 h at 37°C in 50mM ammonium bicarbonate (pH = 7.8). The reactions were terminated by adding trifluoroacetic acid. Samples were concentrated to a final volume of 100 µL using vacuum centrifugation. Peptides were desalted using custom-made stage tips assembled by fixing Empore™ C-18 disks and conditioned with acetonitrile. They were pre-equilibrated with 0.1% formic acid. Acidified samples were subsequently loaded and passed through the stage tips thrice. After washing the stage tips with 0.1% formic acid, the mixture of peptides was eluted using 50% acetonitrile in 0.1% formic acid. For LC-MS analysis, the samples were dried in a vacuum centrifuge and resuspended in 0.1% formic acid.

#### LC-MS analysis

Desalted peptides were separated on an Ultimate 3000 RSLC nano-UPLC system (Thermo Fisher Scientific) connected with an Orbitrap Elite hybrid mass spectrometer. 500 ng of peptides were loaded with automated injections onto a trap column (Acclaim PepMap™ 100, 3 µm particle size, 75 µm × 2 cm) for 5 min, with a flow rate of 5 µl min^−1^ with 0.1% formic acid. The peptides were gradually eluted from the trap column using an increasing concentration of acetonitrile and then separated on C-18 analytical column (PepMap™ RSLC, 2 µm particle size, 100 Å pore size, 75 µm × 50 cm) at 250 nL min^−1^ flow rate. The solvents used for peptide separation were solvent A - 0.1% formic acid in water, and solvent B - 0.1% formic acid in 100% acetonitrile (ACN). After 10 min equilibration, a sharp gradient from 1% to 5% B in 3 mins was run. This was followed by a shallow gradient from 5% to 22% B in 135 mins, which was then extended to 28% B in additional 20 mins. The analytical column temperature was set at 40°C and was directly connected to a nanospray electron ionization source (Thermo Fisher Scientific), which generated a stable spray at a voltage of 1.8 kV. The capillary temperature was set at 275°C for effective nebulization and mobility of peptide ions into the mass spectrometer. The data were acquired in the positive mode using data-dependent acquisition (DDA). Twenty most abundant parent ions from each MS1 spectrum (350-2000 m/z) were sequentially isolated and fragmented by collision-induced dissociation (CID). MS1 and MS2 spectra were acquired in Orbitrap at 60,000 resolution and in ion trap at rapid scan mode, respectively. The maximum filling time for MS1 was set at 100 ms with a limitation of 1×10^6^ ions, whereas for MS2 spectra, the filling time was set at 50 ms with a target ion limitation set at 5×10^3^ ions. Peptide fragmentation was done at a normalized collision energy of 35% with an activation time of 10 ms. Dynamic exclusion of 30 seconds was applied with a repeat count of 1.

#### Protein Identification

The raw files for all the samples were analyzed together by Maxquant (version 2.0.3.0) using the reference proteome of *M. smegmatis mc*^*2*^*155* (https://mycobrowser.epfl.ch/; version v3) using its internal peptide search engine Andromeda. For protein identifications, the following parameters were used: maximum missed cleavages, 2; mass tolerance for first and main search, 20 and 4.5 ppm respectively; mass tolerance for fragment ion, 0.5 Da; variable modifications used were N-terminal acetylation and methionine oxidation; minimum unique peptide required for identification, 1; minimum peptide length for identification, 7; max. peptide mass, 4600 Da; peptide spectrum match identification and protein inference false discovery rate were set at 0.01; dependent peptide and match between run options were enabled with a match time window of 0.7 min and alignment window of 20 min. iBAQ mode was enabled for protein quantification. Mtb orthologues and localization of obtained proteins were identified using Mycobrowser (https://mycobrowser.epfl.ch/) and Uniprot (https://www.uniprot.org/). For proteins whose localization was unknown, PSORTb (https://www.psort.org/psortb/) and Gpos-mPLoc (http://www.csbio.sjtu.edu.cn/bioinf/Gpos-multi/) were used to predict based on amino acid sequences.

## Results

### Synthesis and characterization of AuNP and OMV-AuNP

In order to stabilize OMVs using core solid particles, we first synthesized nanoparticles of diameter ∼50 nm size that can be used to coat with nano-sized OMVs. Gold nanoparticles were the material of choice due to the ease of synthesis, tunable properties, and their ability to enhance immunogenic activity [36-38]. AuNP were synthesized by the one-pot seeded growth method [35]. AuNP seeds were synthesized and used to grow the nanoparticles to the desired size. The synthesis of seeds and nanoparticles were confirmed from the peaks observed in their respective absorption spectra (**Fig 1A**). Further, the size and shape of the particles were characterized using DLS (**Fig 1B**) and TEM (**Fig 1C**). The seeds were 22 ± 3.83 nm in size, and the nanoparticles were 51 ± 2.66 nm in size.

**Figure 1:**
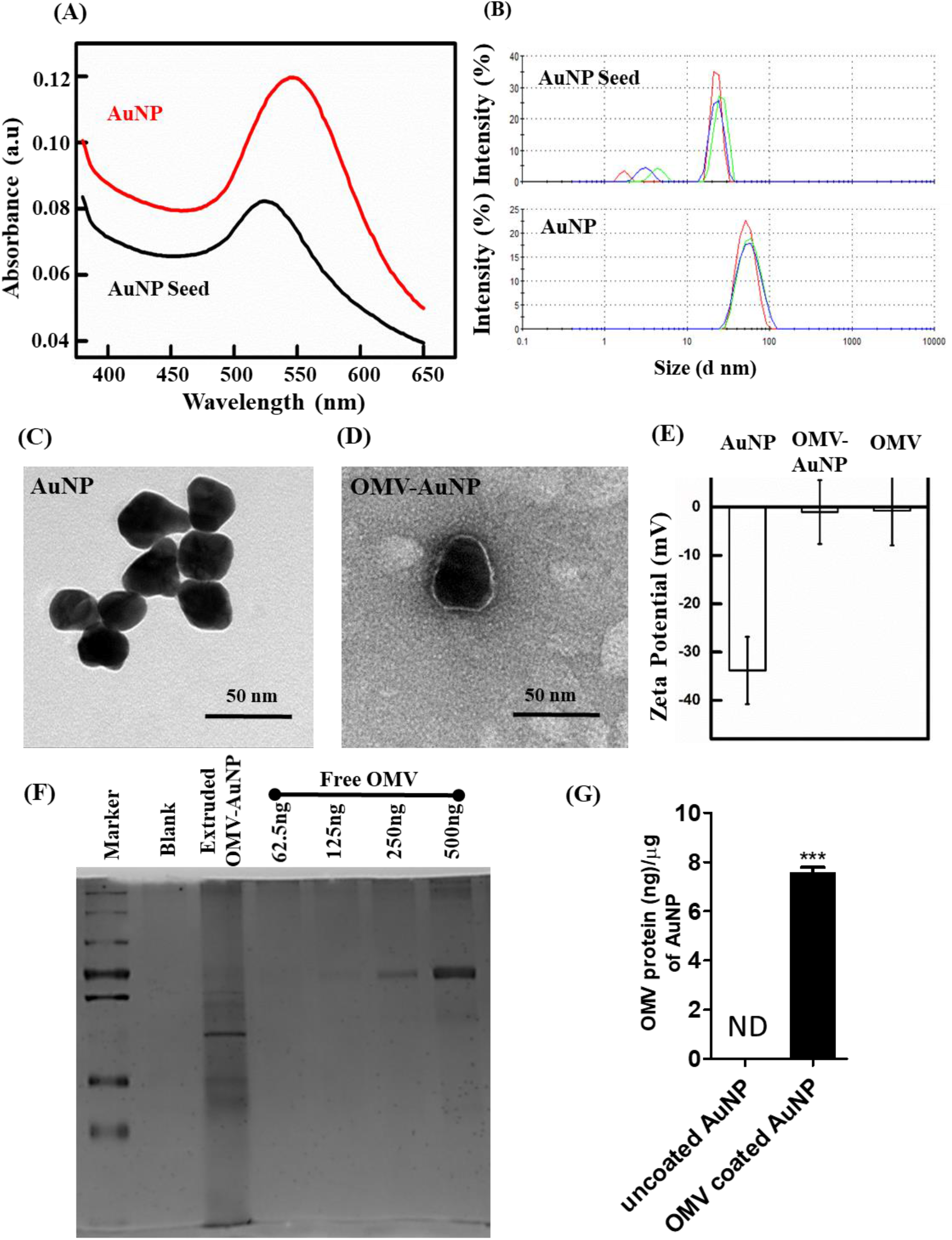
Characterization of synthesized AuNP using one-pot seeded growth method. (**A**) Representative absorption spectra of AuNP seeds and AuNP in water. The peaks at 524 nm and 544 nm confirmed the synthesis of AuNP seeds and AuNP, respectively. (**B**) Size distribution of AuNP seeds and AuNP analyzed by dynamic light scattering. (**C, D**) Representative TEM images of AuNP and OMV-AuNP. Scale bar: 50 nm. (**E**) Zeta potentials of AuNP with and without OMV coating compared to that of free OMVs. (**F**) SDS-PAGE image shows several proteins on OMV-AuNP and various concentrations of free OMVs. (**G**) Quantification of SDS-PAGE protein bands of OMV-AuNP. Data in graphs represent the mean ± s.d.

For membrane coating on AuNP, OMVs were extruded nine times with AuNP through a 100 nm porous membrane to generate OMV-AuNP. Forcing hollow OMVs and AuNP through the membranes ensured coating of AuNPs with OMVs. TEM imaging of OMV-AuNP showed a clear core-shell morphology compared to the uncoated AuNP (**Fig 1C-D, Supplementary Fig 1**). Zeta potential measurements of particles also suggested the presence of the membrane coating as the zeta potential of AuNP increased from -33.8 ± 6.96 mV to - 1.01 ± 6.61 mV after extrusion with OMVs, which was comparable to free OMVs having - 0.719 ± 7.16 mV zeta potential (**Fig 1E**).

SDS-PAGE was performed to determine the protein profile of OMV-AuNP and free OMVs. SDS-PAGE of OMV-AuNP and free OMVs in differing quantities showed the enrichment of several protein bands on OMV-AuNP group compared to free OMVs (**Fig 1F**). A few new bands can be seen in OMV-AuNP compared to free OMVs. This could be due to enrichment of surface proteins in OMV-AuNP as compared to free OMVs for same amount of protein weight (also seen in our Mass-spectrometry data described later). Quantifying the bands of OMVs showed 7 ± 1.11 ng OMV protein coating per microgram of AuNP (**Fig 1G**). The membrane coating was also quantified using the CBQCA assay. Consistent with the SDS-PAGE data, CBQCA quantification measured around 7.6 ± 0.17 ng of OMV coating per microgram of AuNP. A significant drawback of using OMVs is their low stability [32-34]. Hence, we checked the physical stability of synthesized AuNP, isolated OMV, or extruded OMV-AuNP nanoparticles for over three weeks (**Supplementary Fig 2A**). OMVs showed size variation ranging from 147.3 ± 30.78 to 58.29 ± 7.51 nm by day 22, possibly due to aggregation and degradation. However, AuNP and OMV-AuNP showed similar sizes ranging from 36.6 ± 1.56 to 28.49 ± 7.62 nm and 39.11 ± 4.74 to 24.6 ± 3.26 nm, respectively. To check the suspension stability of OMV-AuNP, absorbance readings were taken over time. Transmittance obtained from these measurements showed stable readings (**Supplementary Fig 2B**). Collectively, these results suggest that AuNP coated with OMVs were stable.

**Figure 2:**
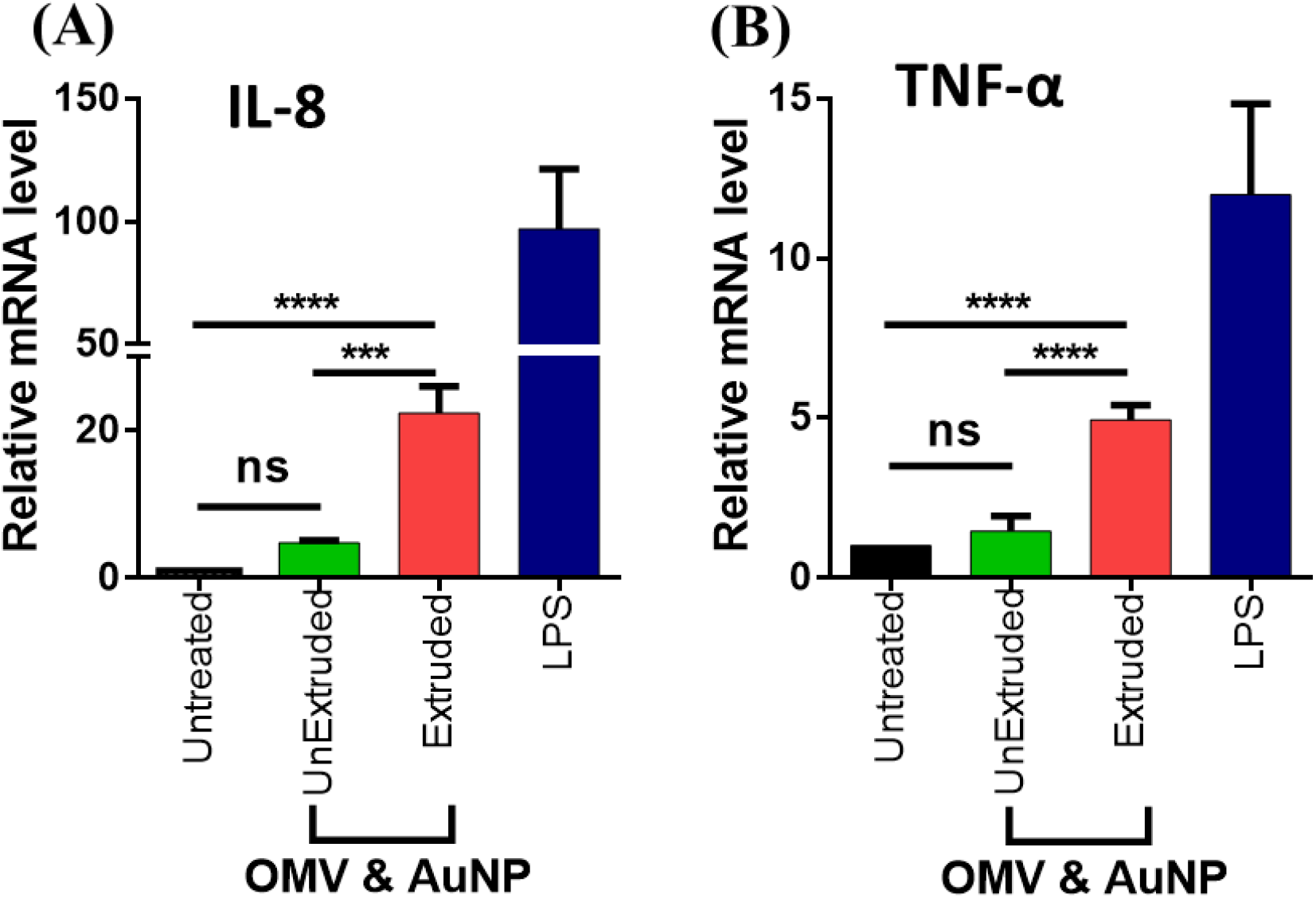
THP1 cells treated with OMV-AuNP showed upregulation of (**A**) hIL-8 and (**B**) hTNF-α mRNA levels compared to their respective controls. Data in the graphs represent the mean ± s.d. ns represents not significant, \*\*\*\**p* < 0.0001 and \*\*\**p* < 0.001. Statistical significance was determined by using ordinary one-way analysis of variance (ANOVA) followed by Tukey’s post-hoc test.

### OMV coated gold nanoparticles elicit a high innate immune response

Bacterial OMVs present a wide array of surface antigens and pathogen-associated molecular patterns (PAMPs) and are capable of generating an immune response. To evaluate whether OMV-AuNP retain these immune-activation properties, we first optimized the ratios of OMVs and AuNP to be used for extrusion to obtain a saturating immune response. As shown in **Supplementary Fig 3**, IL-8 mRNA production in THP-1 cells increased up to 3:1 ratio of OMV:AuNP, after which no further increase in mRNA levels was observed when incubated with 20 µg/mL of gold particles extruded with different amounts of OMVs. Hence, for all future experiments, we used a ratio of 3:1 ratio (protein content on OMVs: gold particles) for extrusion. Next, we tested the effect of increasing OMV-AuNP concentration on immune response by human macrophages. We found that 20 µg/mL of OMV-AuNP (total gold concentration) was the saturating concentration as shown by IL-8 mRNA levels (**Supplementary Fig 4**). All subsequent experiments were done with 20 µg/mL OMV-AuNP.

Remarkably, when cells were treated with free OMVs and OMV-AuNP (equivalent protein content), we observed several-fold higher mRNA levels of hIL-8 (25-fold, **Fig 2A**) and hTNF-α (5-fold, **Fig 2B**) in cells treated with OMV-AuNP compared to media control. Free OMVs (unextruded) also showed a slight increase in mRNA levels but was consistently lower than OMV-AuNP. Subsequent studies carried out at different time points showed that mRNA levels peaked at 24 h (**Supplementary Fig 5**). To determine if this effect was cell line-specific or general, we performed these experiments on primary bone marrow-derived dendritic cells (BMDCs) to gauge the potency of OMV-AuNP to instigate an immune response. In line with the THP-1 data, these cells also exhibited higher levels of mRNA for mIL-6 (**Fig 3A**), mTNF-α (**Fig 3B**), and mIL-1β (**Fig 3C**) when compared to mRNA levels in untreated cells and cells treated with unextruded particles validating the effect of OMV-AuNP on cells.

**Figure 3:**
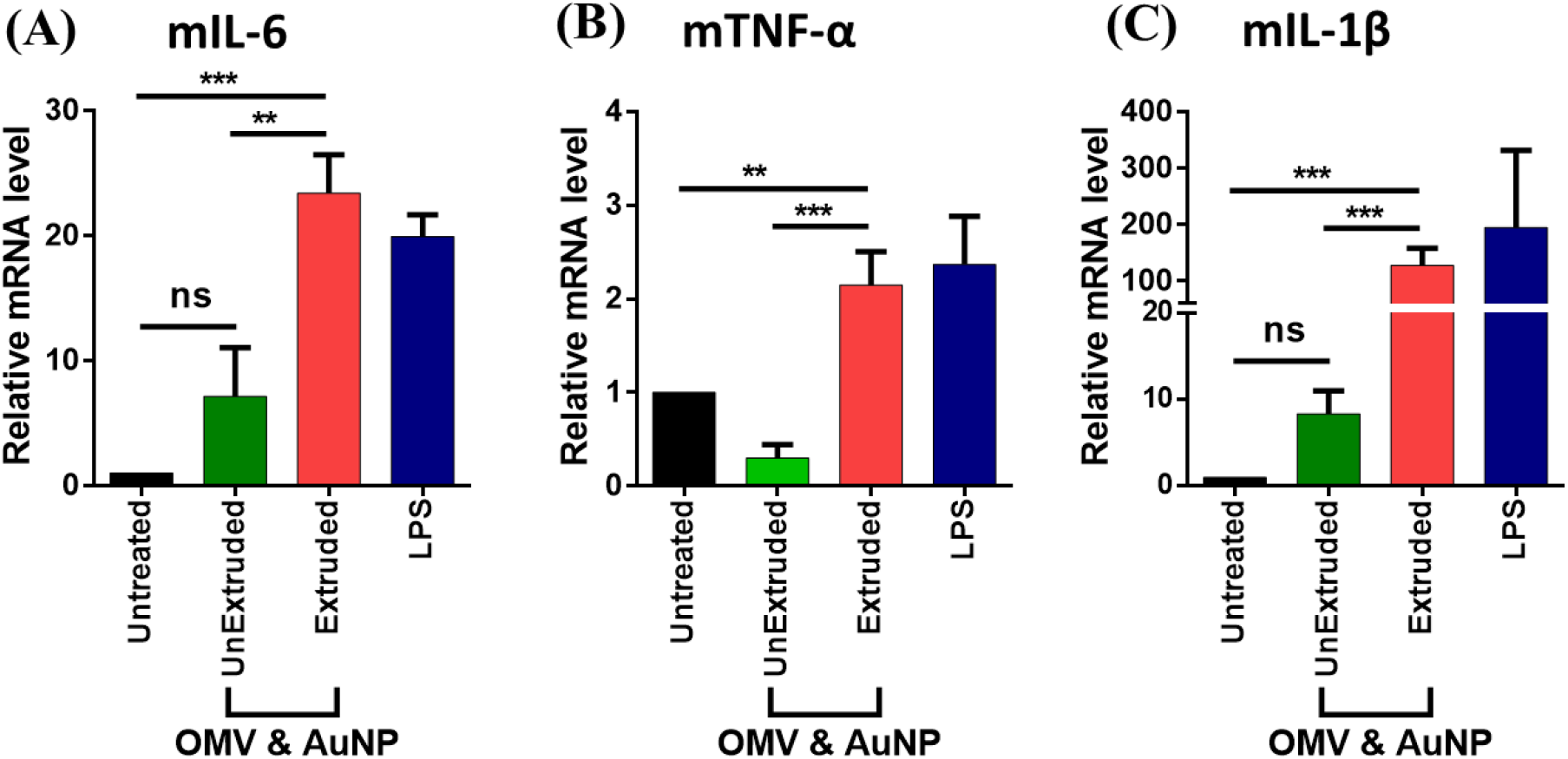
Upregulation of cytokines mRNA level in mouse BMDC cells in response to treatment with extruded OMV-AuNP. Cells treated with OMV-AuNP showed upregulation of (**A**) mIL-6, (**B**) mTNF-α, and (**C**) mIL-1β mRNA levels compared between extruded and unextruded OMVs and AuNP. Data in the graphs represent mean ± s.d. ns represents not significant, \*\*\**p* < 0.001, and \*\**p*<0.01. Statistical significance was determined by using ordinary one-way analysis of variance (ANOVA) followed by Tukey’s post-hoc test.

To further elucidate the effects of OMV-AuNP at a functional level, cell-secreted cytokine proteins were quantified using ELISA. To determine the optimal time to assay for cytokine release, THP-1 cells were treated with OMV-AuNP for various time intervals and levels of hIL-8 in the supernatant were determined. Samples showed a large increase in IL-8 secretion after 24 h and 48 h (**Supplementary Fig 6**). Hence, THP-1 cells were then treated with OMV-AuNP and respective controls for 24 h. The supernatants were collected from these *in vitro* cultures to compare the cytokine signature. Higher levels of hIL-8 (**Fig 4A**) were released by THP-1 cells treated with OMV-AuNP compared to cells treated with AuNP or OMVs alone. Given the co-stimulatory function of hIL-6 in T cell activation, levels of hIL-6 were also quantified. Analysis of the cell supernatants using ELISA showed higher levels of hIL-6 (**Fig 4B**) release from cells treated with OMV-AuNP than cells treated with bare AuNP or OMVs. Similar results were observed when mouse BMDMs were treated with OMV-AuNP. These cells also showed strong mIL-6 (**Fig 4C**) and mTNF-α (**Fig 4D**) release patterns when compared to their respective controls.

**Figure 4:**
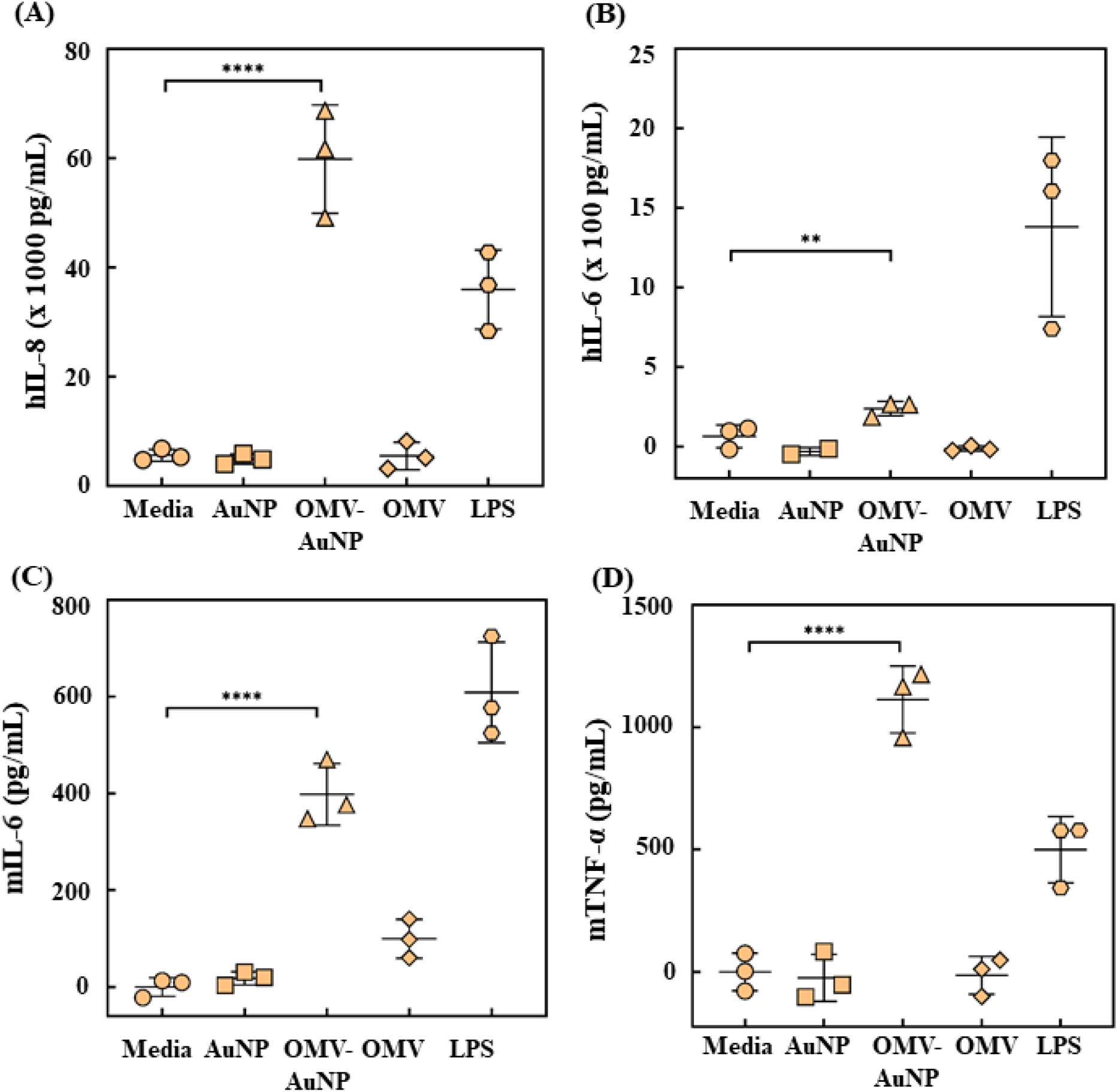
THP-1 cells treated with OMV-AuNP showed significantly increased cytokine release. The concentration of (**A**) hIL-8 and (**B**) hIL-6 cytokines released into the media by THP-1 cells as determined by ELISA. Mouse BMDM cells treated with OMV-AuNP showed pronounced release of cytokines (**C**) mIL-6 and (**D**) mTNF-α in the media as determined by ELISA. Data in the graphs represent the mean ± s.d. \*\*\*\**p* < 0.0001 and \*\**p* < 0.01 were determined by ordinary one-way analysis of variance (ANOVA) followed by Dunnett’s multiple comparison test.

### OMV coated gold nanoparticles show high uptake in THP-1 cells

To understand the mechanism for high immune cell activation, we first evaluated if there was any difference in cell uptake by phagocytic cells such as macrophages among various formulations. Enhanced cell uptake can result in high lysosomal degradation and activation of TLRs present in endosomes and lysosomes [39]. Microscopy studies showed that OMV-AuNP exhibited enhanced particle uptake in THP-1 cells than free OMVs and bare AuNPs when incubated with the same protein and gold content, respectively (**Fig 5A**). The mean fluorescence intensity analysis in the microscopic images showed significantly higher nanoparticle uptake in macrophages treated with OMV-AuNP than free OMVs or unextruded mixture of AuNP and OMVs (**Fig 5B**).

**Figure 5:**
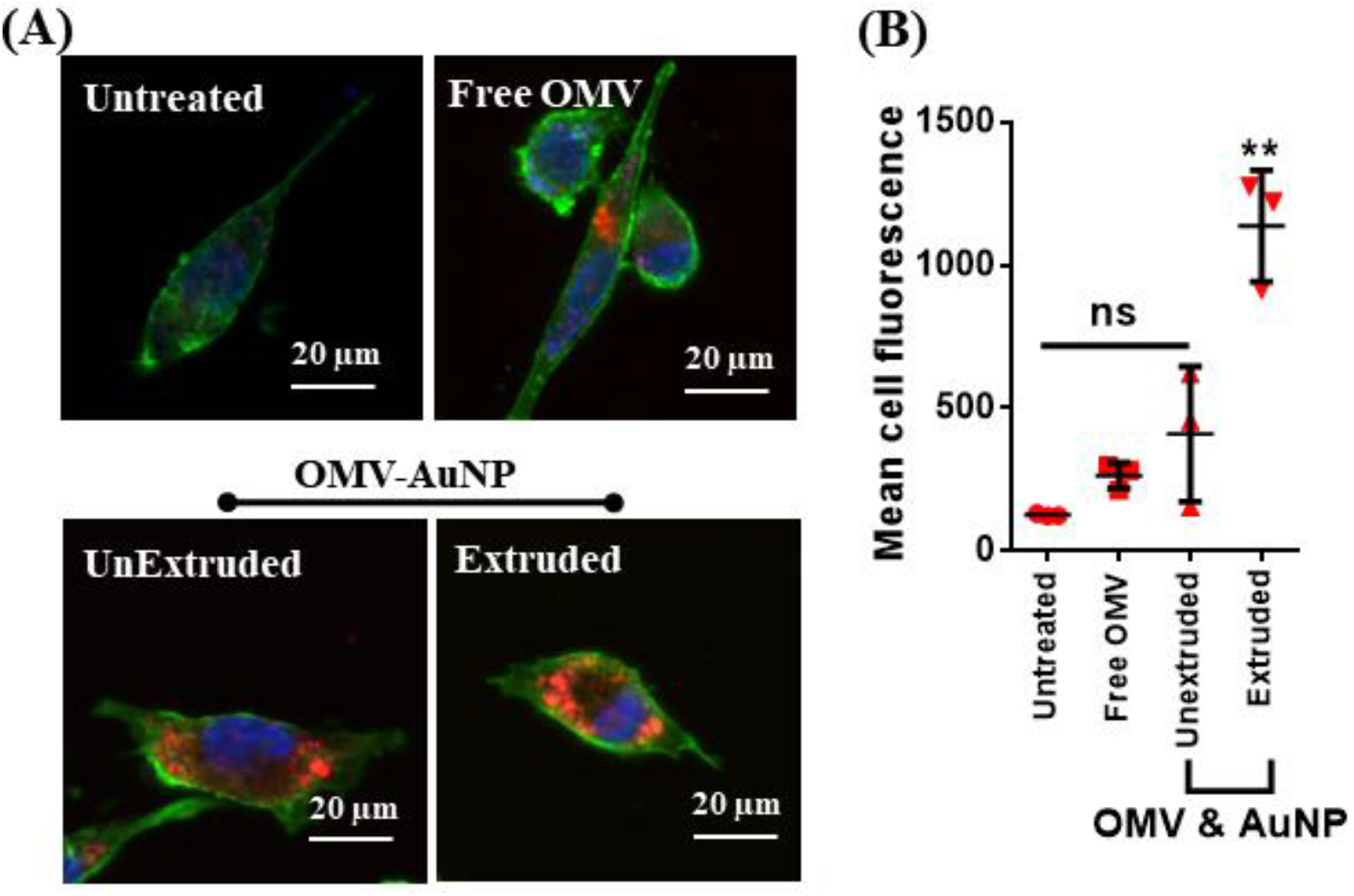
THP-1 cells treated with OMV-AuNP exhibit enhanced cell uptake. (**A**) Representative images of THP-1 cells treated with free OMVs, extruded or unextruded OMV-AuNP while untreated cells served as the control. The OMVs were labeled using DiI dye (red), and the cells were traced using actin (green) and nucleus (blue) staining. (**B**) Quantification of the fluorescence analyzed from labeled OMVs. Data in the graphs represent mean ± s.d., \*\**p* < 0.01 versus all groups. Statistical significance was determined by using ordinary one-way analysis of variance (ANOVA) followed by Tukey’s post-hoc test.

### TLR 2 and TLR 4 are involved in the immune response to OMV-AuNP

Since both proteins and lipids are known to activate the immune response, we wanted to examine the major component of OMVs that results in immune activation. Therefore, extruded OMV-AuNP were treated with proteinase K to degrade exposed proteins (**Fig 6A**). First, the activity of proteinase K was examined by treating free OMVs for 30 min. The absence of several protein bands in the enzyme-treated OMVs compared to untreated OMVs indicates that proteinase K effectively degrades exposed proteins (**Fig 6B**). Next, the extruded OMV-AuNP particles were treated with proteinase K. Cell studies showed significantly decreased mRNA levels for cytokines hIL-8 (8-fold lower, **Fig 6C**) and hTNF-α (3-fold lower, **Fig 6D**) in cells incubated with proteinase K treated OMV-AuNP than those with untreated OMV-AuNP. Together, these studies show that the proteins in OMV-AuNP play an essential role in determining the cell response.

**Figure 6:**
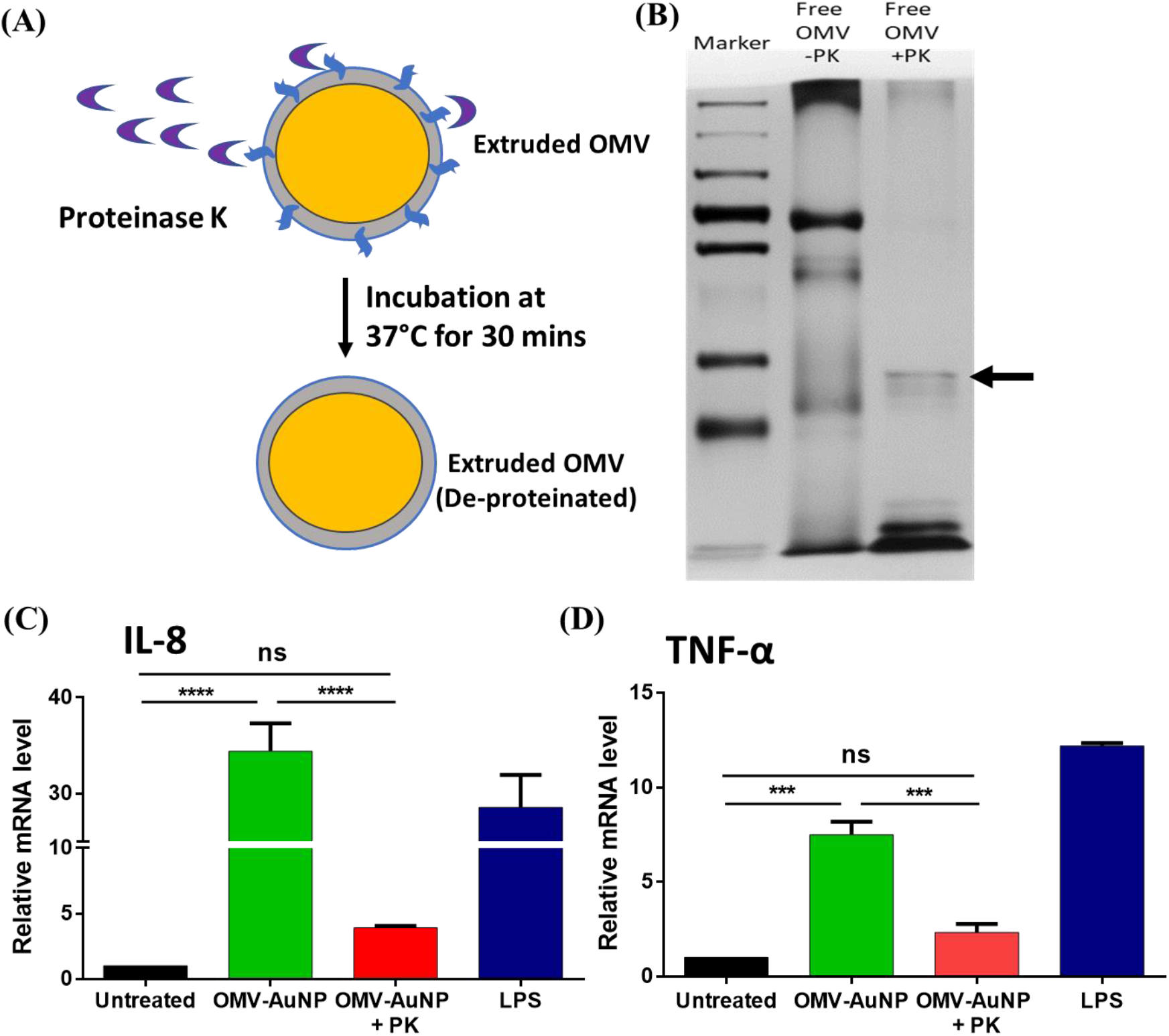
Intact OMV proteins contributed to higher mRNA levels for hIL-8 and hTNF-α cytokines. (**A**) Schematic representation of proteinase K activity on OMV-AuNP. (**B**) SDS-PAGE gel showing bands from OMVs treated in the presence or absence of proteinase K. Arrow indicates proteinase K band. Cells treated with OMV-AuNP showed pronounced mRNA levels for cytokines (**C**) hIL-8 and (**D**) hTNF-α, when the OMVs were intact when compared to proteinase K (PK) treated OMVs. Data in the graphs represent the mean ± s.d. ns represents not significant, \*\*\*\**p* < 0.0001 and \*\*\**p* < 0.001. Statistical significance was determined by using ordinary one-way analysis of variance (ANOVA) followed by Tukey’s post-hoc test.

Several studies in the past have established TLRs’ role in mediating immune response in the presence of OMVs [40, 41]. Lipoproteins and other PAMPs on OMVs can I nteract with TLR2 and TLR4 on antigen-presenting cells [20]. Hence, sparstolonin B (SsnB), a selective TLR antagonist for TLR2 and TLR4 [42], was used to better understand the signaling involved during OMV-AuNP and cell interaction. Initial studies with SsnB on THP-1 cells were focused on optimizing the concentration to alter the immune response due to LPS. The mRNA levels for cytokines hIL-8 (**Supplementary Fig 7A**) and hTNF-α (**Supplementary Fig 7B**) were downregulated in LPS-exposed cells treated with SsnB, which is indicative of the blocking of TLRs on the cells. On optimizing the SsnB to 50 µM, studies on THP-1 cells were performed in the presence of OMV-AuNP. Treating the cells with SsnB in the presence of OMV extruded particles significantly reduced mRNA levels of hTNF-α (**Fig 7A**) and hIL-8 (**Fig 7B**) compared to the no-inhibitor treated group. This shows that TLR2 and TLR4 are the primary signaling mechanism responsible for immune response with OMV-AuNP.

**Figure 7:**
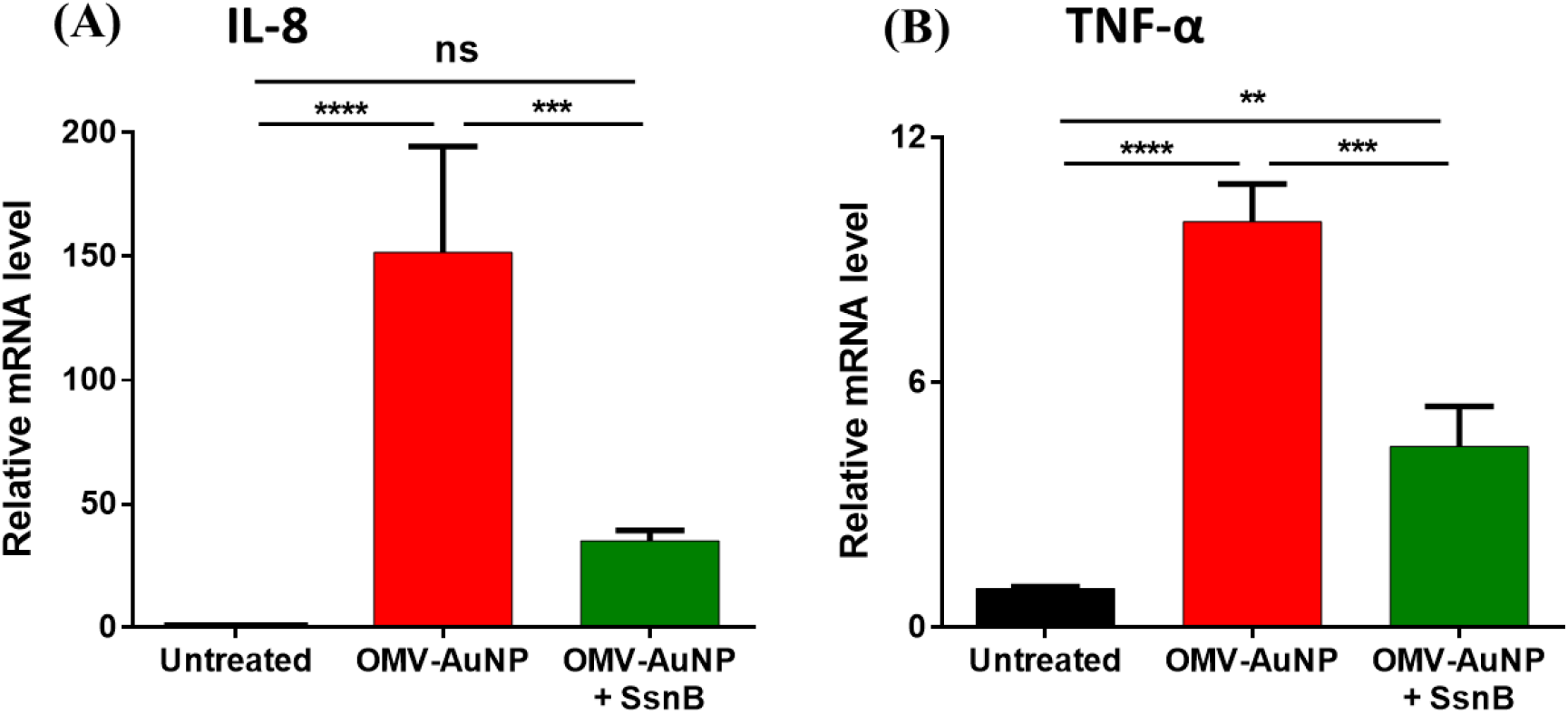
mRNA levels for cytokine production by THP-1 cells treated with OMV-AuNP were mediated by TLR 2 and 4. Downregulation of mRNA level of (**A**) hIL-8 and (**B**) hTNF-α cytokines in response to OMV-AuNP treatment when TLR 2 and 4 were blocked using a TLR agonist. Data in the graphs represent the mean ± s.d. ns represents not significant, \*\*\*\**p* < 0.0001, \*\*\**p* < 0.001, and \*\**p* < 0.01. Statistical significance was determined by using ordinary one-way analysis of variance (ANOVA) followed by Tukey’s post-hoc test.

### Identification of OMV associated proteins via LC-MS/MS analysis

Since proteinase-K treatment of extruded OMV-AuNP particles resulted in reduced immune response as confirmed by lower levels of mRNAs for cytokines hIL-8 and hTNF-α, we next sought to decipher the protein composition of OMVs. We performed an untargeted LC-MS/MS analysis of proteins derived from free OMVs and OMV-AuNP. Three independent samples were prepared, each for free OMVs and OMV-AuNP. Tandem mass-spectrometry analysis identified a total of 43 proteins, with both peptide (PSM) and protein identification FDR set at 0.01. Proteins identified with at least one unique peptide were included for the subsequent analysis. The complete list of identified proteins is given in **Supplementary Table 2**.

To gain insight into the biology of OMV associated proteins, we assigned functional category information to each protein from Mycobrowser either directly or derived from Uniprot and their *M. tuberculosis* orthologues. Out of 43 proteins, 18 proteins have orthologues in *M. tuberculosis*. Functional classification **(Fig 8A)** showed enrichment of three major functional categories: intermediary metabolism and respiration (30.2%), conserved hypotheticals (23.2%), cell wall and cell processes (16.2%), which is in line with a previous report on OMV associated proteins of H37Rv [43]. Among the cell wall and cell processes category, MmpL5 protein and several ABC transporters were prevalent, which have been previously shown to be surface exposed and immunogenic [44]. For instance, PstS, a phosphate transporter of *M. tuberculosis*, was highly immunogenic and conferred protection against intravenous *M. tuberculosis* exposure [45, 46]. In addition to cell envelope-associated proteins, some cytosolic proteins related to metabolism and 10 kDa GroES chaperon were also detected in OMVs. It remains to be investigated if their presence in OMVs is due to their possible interaction with membrane components or they are selectively packaged and secreted via OMVs for specific functions.

**Figure 8:**
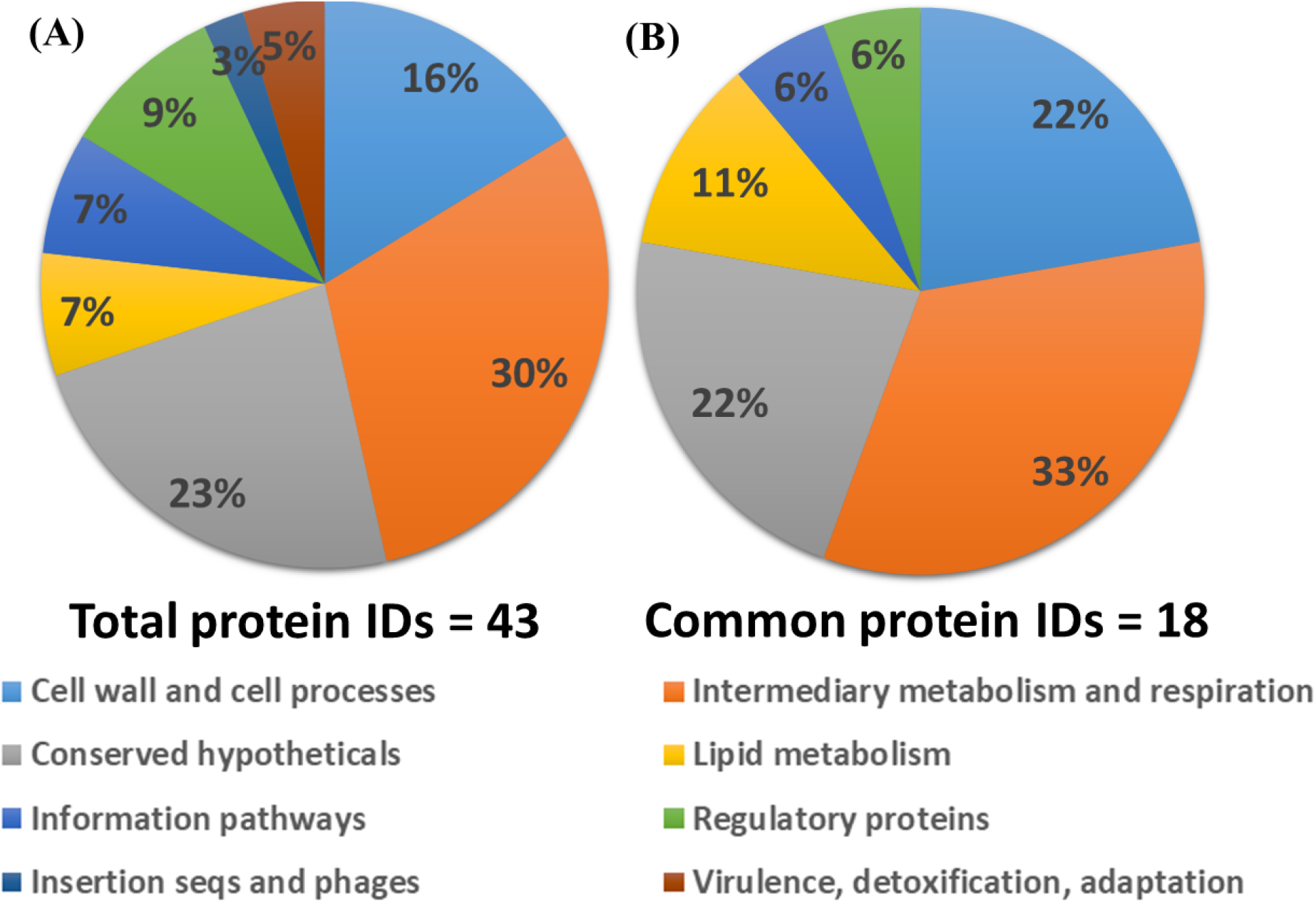
Functional classification of (**A**) total identified proteins and (**B**) proteins commonly present on free OMVs and OMV-AuNPs, based on Mycobrower functional category information.

Next, we wanted to identify the proteins from free OMVs that were also coated on AuNP. First, from the total list of 43, we confirmed the presence of 40 proteins in free OMV samples, and similarly, 21 proteins were confirmed to be present in OMV-AuNP samples. We then performed a comparative analysis to identify the proteins present in both conditions. A total of 18 proteins were identified in both conditions listed in detail in **Table 1**. As expected, most of the OMV-AuNP associated proteins are a subset of free-OMV proteins, except for the three proteins, where one is an ABC transporter and two others are of unknown functions.

Of these 18 retained proteins on OMV-AuNP, eight proteins have orthologues in *M. tuberculosis* H37Rv, suggesting the possibility of generating cross-reactive TB immunity using OMV-AuNP. Similar functional classification analysis **(Fig 8B)** showed proteins belonging to intermediary metabolism and respiration (33.3%), conserved hypotheticals (22.2%), cell wall, and cell processes (22.2%) to be present in higher frequency in OMV-AnNPs. Few of the proteins have unknown functions and cellular localization, thus the exact reason for their presence and possible immune-modulatory roles remains to be understood. On the other hand, proteins belonging to lipid metabolism, regulatory proteins, and information pathways were detected at lower frequencies. Interestingly, MSMEG_3618 (Rv1860) has been reported to be a T-cell reactive immunogenic protein [47]. It remains to be checked which of these OMV-AuNP associated proteins are highly immunogenic and can confer cross-reactive immunity against *M. tuberculosis*. Together, in this study, we identify the protein composition of *M. smegmatis* OMVs and show that the OMV associated proteins can be coated on gold nanoparticles. These OMV-AuNPs have higher uptake in macrophages show enhanced immune response via TLR-2 and TLR-4.

## Discussion

The inability of BCG to prevent *M. tuberculosis* infection in adults is a significant concern towards containing or eradicating TB. This necessitates the development of new vaccines strategies [48]. This study explored the immunomodulating potential of mycobacterial outer membrane vesicle-coated gold nanoparticles. OMVs contain a large number of protein and lipid antigens and could help elicit a broad and robust immune response [21-24]. Our data showed that bio-functionalizing gold nanoparticles by coating OMVs were capable of generating immune response as observed by the cytokine profile. This highlights the possible potential of OMV coated gold nanoparticles in taking us one step closer to combating TB.

Exosomes derived from *Mycobacterium bovis* and Mtb infected macrophages were shown to contain mycobacterial lipid components that can activate macrophages and antigen-specific CD4 and CD8 T cells [49, 50]. Conversely, it was also shown that bacterial OMVs and exosomes from Mtb infected macrophages inhibit T-cell activation due to the presence of immunosuppressive lipoglycans and lipo-proteins such as lipoarabinomannan [51]. *M. tuberculosis* lipids are also known to prevent phagosomal maturation [52-54]. Hence, we focused on using the OMVs derived from *M. smegmatis* which are not known to prevent phagosomal maturation. OMVs from *H. pylori* [55], *P aeruginosa* [56], and *P. gingivalis* [19] have been shown to stimulate macrophages and dendritic cells. However, in case of *M. smegmatis* OMVs, the activation of macrophages was not very high. A key difference is the absence of LPS in Mycobacterium species that can act as a powerful stimulant and might explain the lack of activation with free OMVs in this study. Similar results had been previously reported that showed the lack of immune cell activation with *M. smegmatis* OMVs [43].

The main challenges in using OMVs for vaccine development are their heterogeneous size and composition, fragile nature, and low stability in biological buffers. Hence, for the better delivery of vaccines and enhanced immune response, we combined OMVs with nanoparticles [57]. Nanoparticles are a promising platform for developing vaccines given the high surface area to mass ratio and tunable physical properties. Additionally, nanoparticles improve antigen stability and immunogenicity, which are imperative for designing and developing vaccines [58]. Gold nanoparticles, being non-toxic and biocompatible, have garnered a lot of research interest in targeted delivery for several biomedical applications. Several studies have utilized AuNP for drug and antigen delivery in cancer and viral diseases [34]. In this study, AuNP were used as the core nanoparticle for delivering and presenting OMVs to antigen-presenting cells. Since the size of OMVs are <250 nm, we synthesized 50 nm gold nanoparticles to effectively coat them with OMVs. Gold nanoparticles bio-functionalized with nanogram levels of *M. smegmatis* OMVs resulted in robust activation and cytokine production from macrophages and dendritic cells. The exact mechanism of how gold stabilized OMVs enhance immune response is unclear. We found that OMV-AuNP had higher uptake compared to free OMVs, which could enhance the engagement of intracellular TLRs. Furthermore, treatment of proteinase K resulted in a substantial decrease in cytokine production, suggesting that the major activating moieties are proteins and glycoproteins. Finally, the use of TLR2 and TLR4 inhibitors also decreased cytokine production, linking these two TLRs as a major player in the immune response to OMV-AuNP. TLR2 exists as a heterodimer with other TLRs, such as TLR1 and TLR6, and it is not clear whether these TLRs are also involved in the activation observed with OMV-AuNP [59]. More studies are required to understand this effect.

OMV biogenesis and subsequent release into the extracellular environment of the microbe are initiated by several factors that include oxidative stress [60], altered temperature [61], and iron depletion [62]. Therefore, the production of OMVs in an *in vitro* system can be enhanced by altering these external stimuli or by genetic manipulation of parent bacteria, resulting in designer OMVs, that can be isolated and purified to elicit an immune response or explore therapeutic avenues. The use of OMVs would also expose new and diverse antigens that can result in antibody-based immunity against TB, an important player and present in uninfected healthcare workers who are heavily exposed to TB [45, 63-67].

The efficacy of vaccines depends on the immunogenicity of the vaccine. In this regard, OMVs seem to be an ideal candidate owing to their suitable size range [68] and membrane composition [69]. Our results suggest that THP-1 cells treated with OMVs release several cytokines, including IL-8, a chemokine responsible for recruiting T lymphocytes. Earlier reports have also shown the role of OMVs in activating antigen-presenting cells and inducing specific T cell activation, making them attractive vaccine candidates [18]. The **Error! Bookmark not defined**.vaccine’s design in this study combines the merits of two materials, OMVs and AuNP, to elicit a favorable immune response. Extruding AuNP with OMVs allows the coating of AuNP with suitable antigenic properties that imitate the bacterial pathogen, allowing them to better engage with immune cells. Additionally, OMV-AuNP maintained uniform particle size, which is a drawback with free OMVs.

There are a few limitations of this study. We have used gold particles, which are non-degradable and can accumulate in the body. The use of degradable polymeric particles could overcome this limitation. All experiments were done *in vitro* with antigen-presenting cells, and this platform needs to be further evaluated for induction of adaptive immunity and *in vivo* efficacy. Finally, OMVs exhibit high batch-to-batch variability, which is expected to be reflected on OMV-AuNP as well. We did observe slight changes in the magnitude of activation from different batches of OMVs; however, all batches consistently showed activation. Nonetheless, OMVs are already in clinical use for *Neisseria meningitidis* and have shown efficacy against serogroup B meningococcal disease [25], which shows that batch variability can be overcome and is not a significant limitation for clinical translation. In summary, we have demonstrated that mycobacterial outer membrane vesicle-coated gold nanoparticles can elicit a high immune response under *in vitro* conditions and overcome challenges associated with free OMVs. The platform offers an alternative strategy for the development of TB vaccines.

## Acknowledgements

We thank the Central Facilities at Centre for BioSystems Science and Engineering, Materials Engineering, Molecular Reproduction, Development and Genetics (MRDG), Molecular Biophysics Unit (MBU), and Microbiology and Cell Biology (MCB) for providing access to instruments. We also thank the Central Animal Facility at the Indian Institute of Science for breeding and maintaining mice. We are thankful to Prof. Sandhya S. Visweswariah for providing access to use the DLS and Ultracentrifuge. We acknowledge DST-FIST for providing access to the TEM facility. Technical discussions with Dr. Siddharth Jhunjhunwala were also helpful in designing experiments.

## Author contributions

Study design and experimental planning: EG, AG, PP, and RA; Investigation: EG, AG, TL, PP, SI, IG, LS, SJ, PRS and AAD; Supervision: RM and RA; Funding acquisition: RM and RA; Writing – original draft: EG and RA. All authors reviewed, edited, and approved the final version to be published.

## References

[1] G.B.D.C.o.D. Collaborators, Global, regional, and national age-sex specific mortality for 264 causes of death, 1980-2016: a systematic analysis for the Global Burden of Disease Study 2016, Lancet, 390 (2017) 1151–1210. 10.1016/S0140-6736(17)32152-9

[2] L.C. Rodrigues, V.K. Diwan, J.G. Wheeler, Protective effect of BCG against tuberculous meningitis and miliary tuberculosis: a meta-analysis, International journal of epidemiology, 22 (1993) 1154–1158. 10.1093/ije/22.6.1154

[3] B.E. Marciano, C.Y. Huang, G. Joshi, N. Rezaei, B.C. Carvalho, Z. Allwood, A. Ikinciogullari, S.M. Reda, A. Gennery, V. Thon, F. Espinosa-Rosales, W. Al-Herz, O. Porras, A. Shcherbina, A. Szaflarska, S. Kilic, J.L. Franco, A.C. Gomez Raccio, P. Roxo, Jr., I. Esteves, N. Galal, A.S. Grumach, S. Al-Tamemi, A. Yildiran, J.C. Orellana, M. Yamada, T. Morio, D. Liberatore, Y. Ohtsuka, Y.L. Lau, R. Nishikomori, C. Torres-Lozano, J.T. Mazzucchelli, M.M. Vilela, F.S. Tavares, L. Cunha, J.A. Pinto, S.E. Espinosa-Padilla, L. Hernandez-Nieto, R.A. Elfeky, T. Ariga, H. Toshio, F. Dogu, F. Cipe, R. Formankova, M.E. Nunez-Nunez, L. Bezrodnik, J.G. Marques, M.I. Pereira, V. Listello, M.A. Slatter, Z. Nademi, D. Kowalczyk, T.A. Fleisher, G. Davies, B. Neven, S.D. Rosenzweig, BCG vaccination in patients with severe combined immunodeficiency: complications, risks, and vaccination policies, J. Allergy Clin. Immunol., 133 (2014) 1134–1141. 10.1016/j.jaci.2014.02.028

[4] A.C. Hesseling, B.J. Marais, R.P. Gie, H.S. Schaaf, P.E. Fine, P. Godfrey-Faussett, N. Beyers, The risk of disseminated Bacille Calmette-Guerin (BCG) disease in HIV-infected children, Vaccine, 25 (2007) 14–18. 10.1016/j.vaccine.2006.07.020

[5] A.C. Hesseling, L.F. Johnson, H. Jaspan, M.F. Cotton, A. Whitelaw, H.S. Schaaf, P.E. Fine, B.S. Eley, B.J. Marais, J. Nuttall, N. Beyers, P. Godfrey-Faussett, Disseminated bacille Calmette-Guerin disease in HIV-infected South African infants, Bull. World Health Organ., 87 (2009) 505–511. 10.2471/blt.08.055657

[6] C. Aagaard, T. Hoang, J. Dietrich, P.J. Cardona, A. Izzo, G. Dolganov, G.K. Schoolnik, J.P. Cassidy, R. Billeskov, P. Andersen, A multistage tuberculosis vaccine that confers efficient protection before and after exposure, Nature medicine, 17 (2011) 189–194. 10.1038/nm.2285

[7] S.H. Kaufmann, M. Gengenbacher, Recombinant live vaccine candidates against tuberculosis, Current opinion in biotechnology, 23 (2012) 900–907. 10.1016/j.copbio.2012.03.007

[8] M.J. Ahsan, Recent advances in the development of vaccines for tuberculosis, Therapeutic advances in vaccines, 3 (2015) 66–75. 10.1177/2051013615593891

[9] C. Wang, J. Lu, W. Du, G. Wang, X. Li, X. Shen, C. Su, L. Yang, B. Chen, J. Wang, M. Xu, Ag85b/ESAT6-CFP10 adjuvanted with aluminum/poly-IC effectively protects guinea pigs from latent mycobacterium tuberculosis infection, Vaccine, (2019). 10.1016/j.vaccine.2019.06.078

[10] L.K. Schrager, J. Vekemens, N. Drager, D.M. Lewinsohn, O.F. Olesen, The status of tuberculosis vaccine development, Lancet Infect. Dis., (2020). 10.1016/S1473-3099(19)30625-5

[11] M.D. Tameris, M. Hatherill, B.S. Landry, T.J. Scriba, M.A. Snowden, S. Lockhart, J.E. Shea, J.B. McClain, G.D. Hussey, W.A. Hanekom, H. Mahomed, H. McShane, M.A.T.S. Team, Safety and efficacy of MVA85A, a new tuberculosis vaccine, in infants previously vaccinated with BCG: a randomised, placebo-controlled phase 2b trial, Lancet, 381 (2013) 1021–1028. 10.1016/S0140-6736(13)60177-4

[12] H. Ning, W. Zhang, J. Kang, T. Ding, X. Liang, Y. Lu, C. Guo, W. Sun, H. Wang, Y. Bai, L. Shen, Subunit Vaccine ESAT-6:c-di-AMP Delivered by Intranasal Route Elicits Immune Responses and Protects Against Mycobacterium tuberculosis Infection, Front Cell Infect Microbiol, 11 (2021) 647220. 10.3389/fcimb.2021.647220

[13] M. Li, H. Zhou, C. Yang, Y. Wu, X. Zhou, H. Liu, Y. Wang, Bacterial outer membrane vesicles as a platform for biomedical applications: An update, J. Control. Release, 323 (2020) 253–268.https://doi.org/10.1016/j.jconrel.2020.04.031

[14] M. Kaparakis-Liaskos, R.L. Ferrero, Immune modulation by bacterial outer membrane vesicles, Nat. Rev. Immunol., 15 (2015) 375–387. 10.1038/nri3837

[15] S.N. Wai, B. Lindmark, T. Soderblom, A. Takade, M. Westermark, J. Oscarsson, J. Jass, A. Richter-Dahlfors, Y. Mizunoe, B.E. Uhlin, Vesicle-mediated export and assembly of pore-forming oligomers of the enterobacterial ClyA cytotoxin, Cell, 115 (2003) 25–35. 10.1016/s0092-8674(03)00754-2

[16] D.J. Chen, N. Osterrieder, S.M. Metzger, E. Buckles, A.M. Doody, M.P. DeLisa, D. Putnam, Delivery of foreign antigens by engineered outer membrane vesicle vaccines, Proc. Natl. Acad. Sci. U. S. A., 107 (2010) 3099–3104. 10.1073/pnas.0805532107

[17] C.C. Allison, T.A. Kufer, E. Kremmer, M. Kaparakis, R.L. Ferrero, Helicobacter pylori induces MAPK phosphorylation and AP-1 activation via a NOD1-dependent mechanism, Journal of immunology, 183 (2009) 8099–8109. 10.4049/jimmunol.0900664

[18] R.C. Alaniz, B.L. Deatherage, J.C. Lara, B.T. Cookson, Membrane vesicles are immunogenic facsimiles of Salmonella typhimurium that potently activate dendritic cells, prime B and T cell responses, and stimulate protective immunity in vivo, J. Immunol., 179 (2007) 7692–7701. 10.4049/jimmunol.179.11.7692

[19] R. Nakao, H. Hasegawa, K. Ochiai, S. Takashiba, A. Ainai, M. Ohnishi, H. Watanabe, H. Senpuku, Outer membrane vesicles of Porphyromonas gingivalis elicit a mucosal immune response, PLoS One, 6 (2011) e26163. 10.1371/journal.pone.0026163

[20] X. Cao, Self-regulation and cross-regulation of pattern-recognition receptor signalling in health and disease, Nat. Rev. Immunol., 16 (2016) 35–50. 10.1038/nri.2015.8

[21] P. Ogongo, A.J. Steyn, F. Karim, K.J. Dullabh, I. Awala, R. Madansein, A. Leslie, S.M. Behar, Differential skewing of donor-unrestricted and gammadelta T cell repertoires in tuberculosis-infected human lungs, J. Clin. Invest., 130 (2020) 214–230. 10.1172/JCI130711

[22] L. Shen, J. Frencher, D. Huang, W. Wang, E. Yang, C.Y. Chen, Z. Zhang, R. Wang, A. Qaqish, M.H. Larsen, H. Shen, S.A. Porcelli, W.R. Jacobs, Jr., Z.W. Chen, Immunization of Vgamma2Vdelta2 T cells programs sustained effector memory responses that control tuberculosis in nonhuman primates, Proc. Natl. Acad. Sci. U. S. A., 116 (2019) 6371–6378. 10.1073/pnas.1811380116

[23] E. Ribi, R.L. Anacker, W.R. Barclay, W. Brehmer, S.C. Harris, W.R. Leif, J. Simmons, Efficacy of mycobacterial cell walls as a vaccine against airborne tuberculosis in the Rheusus monkey, J. Infect. Dis., 123 (1971) 527–538. 10.1093/infdis/123.5.527

[24] H. Li, L. Liu, W.J. Zhang, X. Zhang, J. Zheng, L. Li, X. Zhu, Q. Yang, M. Zhang, H. Liu, X. Chen, Q. Jin, Analysis of the Antigenic Properties of Membrane Proteins of Mycobacterium tuberculosis, Sci. Rep., 9 (2019) 3042. 10.1038/s41598-019-39402-z

[25] J. Holst, D. Martin, R. Arnold, C.C. Huergo, P. Oster, J. O’Hallahan, E. Rosenqvist, Properties and clinical performance of vaccines containing outer membrane vesicles from Neisseria meningitidis, Vaccine, 27 Suppl 2 (2009) B3–12. 10.1016/j.vaccine.2009.04.071

[26] C.C. Peeters, H.C. Rumke, L.C. Sundermann, E.M. Rouppe van der Voort, J. Meulenbelt, M. Schuller, A.J. Kuipers, P. van der Ley, J.T. Poolman, Phase I clinical trial with a hexavalent PorA containing meningococcal outer membrane vesicle vaccine, Vaccine, 14 (1996) 1009–1015. 10.1016/0264-410x(96)00001-1

[27] I. Claassen, J. Meylis, P. van der Ley, C. Peeters, H. Brons, J. Robert, D. Borsboom, A. van der Ark, I. van Straaten, P. Roholl, B. Kuipers, J. Poolman, Production, characterization and control of a Neisseria meningitidis hexavalent class 1 outer membrane protein containing vesicle vaccine, Vaccine, 14 (1996) 1001–1008. 10.1016/0264-410x(96)00020-5

[28] Y. Tirado, A. Puig, N. Alvarez, R. Borrero, A. Aguilar, F. Camacho, F. Reyes, S. Fernandez, J.L. Perez, R. Acevedo, D. Mata Espinoza, J.A. Payan, M.L. Garcia, R. Kadir, M.E. Sarmiento, R. Hernandez-Pando, M.N. Norazmi, A. Acosta, Mycobacterium smegmatis proteoliposome induce protection in a murine progressive pulmonary tuberculosis model, Tuberculosis (Edinb), 101 (2016) 44–48. 10.1016/j.tube.2016.07.017

[29] L. Rodriguez, Y. Tirado, F. Reyes, A. Puig, R. Kadir, R. Borrero, S. Fernandez, G. Reyes, N. Alvarez, M.A. Garcia, M.E. Sarmiento, M.N. Norazmi, J.L. Perez Quinoy, A. Acosta, Proteoliposomes from Mycobacterium smegmatis induce immune cross-reactivity against Mycobacterium tuberculosis antigens in mice, Vaccine, 29 (2011) 6236–6241. 10.1016/j.vaccine.2011.06.077

[30] Y. Tirado, A. Puig, N. Alvarez, R. Borrero, A. Aguilar, F. Camacho, F. Reyes, S. Fernandez, J.L. Perez, D.M. Espinoza, J.A. Payan, M.E. Sarmiento, M.N. Norazmi, R. Hernandez-Pando, A. Acosta, Protective capacity of proteoliposomes from Mycobacterium bovis BCG in a mouse model of tuberculosis, Human vaccines & immunotherapeutics, 11 (2015) 657–661. 10.1080/21645515.2015.1011566

[31] F. Reyes, Y. Tirado, A. Puig, R. Borrero, G. Reyes, S. Fernandez, J.L. Perez, R. Kadir, C. Zayas, M.N. Norazmi, M.E. Sarmiento, A. Acosta, Immunogenicity and cross-reactivity against Mycobacterium tuberculosis of proteoliposomes derived from Mycobacterium bovis BCG, BMC Immunol., 14 Suppl 1 (2013) S7. 10.1186/1471-2172-14-S1-S7

[32] M.S. Leo van der Pol, and Peter van der Ley, Outer membrane vesicles as platform vaccine technology, Biotechnol J., 10 (2015) 1689–1706.

[33] G. Wu, H. Ji, X. Guo, Y. Li, T. Ren, H. Dong, J. Liu, Y. Liu, X. Shi, B. He, Nanoparticle reinforced bacterial outer-membrane vesicles effectively prevent fatal infection of carbapenem-resistant Klebsiella pneumoniae, Nanomedicine, 24 (2020) 102148. 10.1016/j.nano.2019.102148

[34] W. Gao, R.H. Fang, S. Thamphiwatana, B.T. Luk, J. Li, P. Angsantikul, Q. Zhang, C.M. Hu, L. Zhang, Modulating antibacterial immunity via bacterial membrane-coated nanoparticles, Nano Lett., 15 (2015) 1403–1409. 10.1021/nl504798g

[35] P. Zhang, Y. Li, D. Wang, H. Xia, High-Yield Production of Uniform Gold Nanoparticles with Sizes from 31 to 577 nm via One-Pot Seeded Growth and Size-Dependent SERS Property, Particle & Particle Systems Characterization, 33 (2016) 924–932. 10.1002/ppsc.201600188

[36] E.C. Dreaden, A.M. Alkilany, X. Huang, C.J. Murphy, M.A. El-Sayed, The golden age: gold nanoparticles for biomedicine, Chem. Soc. Rev., 41 (2012) 2740–2779. 10.1039/c1cs15237h

[37] L.A. Dykman, N.G. Khlebtsov, Immunological properties of gold nanoparticles, Chem. Sci., 8 (2017) 1719–1735. 10.1039/c6sc03631g

[38] L.A. Dykman, Gold nanoparticles for preparation of antibodies and vaccines against infectious diseases, Expert Rev Vaccines, 19 (2020) 465–477. 10.1080/14760584.2020.1758070

[39] A. Chaturvedi, S.K. Pierce, How Location Governs Toll-Like Receptor Signaling, Traffic, 10 (2009) 621–628. https://doi.org/10.1111/j.1600-0854.2009.00899.x

[40] N.D. Pecora, A.J. Gehring, D.H. Canaday, W.H. Boom, C.V. Harding, Mycobacterium tuberculosis LprA is a lipoprotein agonist of TLR2 that regulates innate immunity and APC function, Journal of immunology, 177 (2006) 422–429. 10.4049/jimmunol.177.1.422

[41] R. Prados-Rosales, A. Baena, L.R. Martinez, J. Luque-Garcia, R. Kalscheuer, U. Veeraraghavan, C. Camara, J.D. Nosanchuk, G.S. Besra, B. Chen, J. Jimenez, A. Glatman-Freedman, W.R. Jacobs, Jr., S.A. Porcelli, A. Casadevall, Mycobacteria release active membrane vesicles that modulate immune responses in a TLR2-dependent manner in mice, The Journal of clinical investigation, 121 (2011) 1471–1483. 10.1172/JCI44261

[42] Q. Liang, Q. Wu, J. Jiang, J. Duan, C. Wang, M.D. Smith, H. Lu, Q. Wang, P. Nagarkatti, D. Fan, Characterization of sparstolonin B, a Chinese herb-derived compound, as a selective Toll-like receptor antagonist with potent anti-inflammatory properties, The Journal of biological chemistry, 286 (2011) 26470–26479. 10.1074/jbc.M111.227934

[43] R. Prados-Rosales, A. Baena, L.R. Martinez, J. Luque-Garcia, R. Kalscheuer, U. Veeraraghavan, C. Camara, J.D. Nosanchuk, G.S. Besra, B. Chen, J. Jimenez, A. Glatman-Freedman, W.R. Jacobs, S.A. Porcelli, A. Casadevall, Mycobacteria release active membrane vesicles that modulate immune responses in a TLR2-dependent manner in mice, J. Clin. Invest., 121 (2011) 1471–1483. 10.1172/JCI44261

[44] H.S. Garmory, R.W. Titball, ATP-binding cassette transporters are targets for the development of antibacterial vaccines and therapies, Infect. Immun., 72 (2004) 6757–6763. 10.1128/IAI.72.12.6757-6763.2004

[45] A. Watson, H. Li, B. Ma, R. Weiss, D. Bendayan, L. Abramovitz, N. Ben-Shalom, M. Mor, E. Pinko, M. Bar Oz, Z. Wang, F. Du, Y. Lu, J. Rybniker, R. Dahan, H. Huang, D. Barkan, Y. Xiang, B. Javid, N.T. Freund, Human antibodies targeting a Mycobacterium transporter protein mediate protection against tuberculosis, Nat Commun, 12 (2021) 602. 10.1038/s41467-021-20930-0

[46] A. Tanghe, P. Lefevre, O. Denis, S. D’Souza, M. Braibant, E. Lozes, M. Singh, D. Montgomery, J. Content, K. Huygen, Immunogenicity and protective efficacy of tuberculosis DNA vaccines encoding putative phosphate transport receptors, J. Immunol., 162 (1999) 1113–1119.

[47] V. Satchidanandam, N. Kumar, S. Biswas, R.S. Jumani, C. Jain, R. Rani, B. Aggarwal, J. Singh, M.R. Kotnur, A. Sridharan, The Secreted Protein Rv1860 of Mycobacterium tuberculosis Stimulates Human Polyfunctional CD8+ T Cells, Clin. Vaccine Immunol., 23 (2016) 282–293. 10.1128/CVI.00554-15

[48] K. Floyd, P. Glaziou, A. Zumla, M. Raviglione, The global tuberculosis epidemic and progress in care, prevention, and research: an overview in year 3 of the End TB era, The Lancet. Respiratory medicine, 6 (2018) 299–314. 10.1016/S2213-2600(18)30057-2

[49] P.K. Giri, J.S. Schorey, Exosomes Derived from M-Bovis BCG Infected Macrophages Activate Antigen-Specific CD4(+) and CD8(+) T Cells In Vitro and In Vivo, PLoS One, 3 (2008). ARTN e2461 10.1371/journal.pone.0002461

[50] Y. Cheng, J.S. Schorey, Exosomes carrying mycobacterial antigens can protect mice against Mycobacterium tuberculosis infection, Eur. J. Immunol., 43 (2013) 3279–3290. 10.1002/eji.201343727

[51] J.J. Athman, O.J. Sande, S.G. Groft, S.M. Reba, N. Nagy, P.A. Wearsch, E.T. Richardson, R. Rojas, W.H. Boom, S. Shukla, C.V. Harding, Mycobacterium tuberculosis Membrane Vesicles Inhibit T Cell Activation, J. Immunol., 198 (2017) 2028–2037. 10.4049/jimmunol.1601199

[52] K. Sachdeva, M. Goel, M. Sudhakar, M. Mehta, R. Raju, K. Raman, A. Singh, V. Sundaramurthy, Mycobacterium tuberculosis (Mtb) lipid mediated lysosomal rewiring in infected macrophages modulates intracellular Mtb trafficking and survival, J. Biol. Chem., 295 (2020) 9192–9210. 10.1074/jbc.RA120.012809

[53] P.B. Kang, A.K. Azad, J.B. Torrelles, T.M. Kaufman, A. Beharka, E. Tibesar, L.E. DesJardin, L.S. Schlesinger, The human macrophage mannose receptor directs Mycobacterium tuberculosis lipoarabinomannan-mediated phagosome biogenesis, J. Exp. Med., 202 (2005) 987–999. 10.1084/jem.20051239

[54] R.A. Fratti, J. Chua, I. Vergne, V. Deretic, Mycobacterium tuberculosis glycosylated phosphatidylinositol causes phagosome maturation arrest, Proc. Natl. Acad. Sci. U. S. A., 100 (2003) 5437–5442. 10.1073/pnas.0737613100

[55] J. Winter, D. Letley, J. Rhead, J. Atherton, K. Robinson, Helicobacter pylori membrane vesicles stimulate innate pro-and anti-inflammatory responses and induce apoptosis in Jurkat T cells, Infection and immunity, 82 (2014) 1372–1381. 10.1128/IAI.01443-13

[56] X. Zhang, F. Yang, J. Zou, W. Wu, H. Jing, Q. Gou, H. Li, J. Gu, Q. Zou, J. Zhang, Immunization with Pseudomonas aeruginosa outer membrane vesicles stimulates protective immunity in mice, Vaccine, 36 (2018) 1047–1054. 10.1016/j.vaccine.2018.01.034

[57] R. Pati, M. Shevtsov, A. Sonawane, Nanoparticle Vaccines Against Infectious Diseases, Frontiers in immunology, 9 (2018) 2224. 10.3389/fimmu.2018.02224

[58] L. Zhao, A. Seth, N. Wibowo, C.X. Zhao, N. Mitter, C. Yu, A.P. Middelberg, Nanoparticle vaccines, Vaccine, 32 (2014) 327–337. 10.1016/j.vaccine.2013.11.069

[59] L. Oliveira-Nascimento, P. Massari, L.M. Wetzler, The Role of TLR2 in Infection and Immunity, Front. Immunol., 3 (2012) 79. 10.3389/fimmu.2012.00079

[60] A.J. McBroom, M.J. Kuehn, Release of outer membrane vesicles by Gram-negative bacteria is a novel envelope stress response, Molecular microbiology, 63 (2007) 545–558. 10.1111/j.1365-2958.2006.05522.x

[61] T. Baumgarten, S. Sperling, J. Seifert, M. von Bergen, F. Steiniger, L.Y. Wick, H.J. Heipieper, Membrane vesicle formation as a multiple-stress response mechanism enhances Pseudomonas putida DOT-T1E cell surface hydrophobicity and biofilm formation, Applied and environmental microbiology, 78 (2012) 6217–6224. 10.1128/AEM.01525-12

[62] T. Olczak, H. Wojtowicz, J. Ciuraszkiewicz, M. Olczak, Species specificity, surface exposure, protein expression, immunogenicity, and participation in biofilm formation of Porphyromonas gingivalis HmuY, BMC microbiology, 10 (2010) 134. 10.1186/1471-2180-10-134

[63] A. Williams, R. Reljic, I. Naylor, S.O. Clark, G. Falero-Diaz, M. Singh, S. Challacombe, P.D. Marsh, J. Ivanyi, Passive protection with immunoglobulin A antibodies against tuberculous early infection of the lungs, Immunology, 111 (2004) 328–333. 10.1111/j.1365-2567.2004.01809.x

[64] N. Zimmermann, V. Thormann, B. Hu, A.B. Kohler, A. Imai-Matsushima, C. Locht, E. Arnett, L.S. Schlesinger, T. Zoller, M. Schurmann, S.H. Kaufmann, H. Wardemann, Human isotype-dependent inhibitory antibody responses against Mycobacterium tuberculosis, EMBO Mol. Med., 8 (2016) 1325–1339. 10.15252/emmm.201606330

[65] L.L. Lu, A.W. Chung, T.R. Rosebrock, M. Ghebremichael, W.H. Yu, P.S. Grace, M.K. Schoen, F. Tafesse, C. Martin, V. Leung, A.E. Mahan, M. Sips, M.P. Kumar, J. Tedesco, H. Robinson, E. Tkachenko, M. Draghi, K.J. Freedberg, H. Streeck, T.J. Suscovich, D.A. Lauffenburger, B.I. Restrepo, C. Day, S.M. Fortune, G. Alter, A Functional Role for Antibodies in Tuberculosis, Cell, 167 (2016) 433–443 e414. 10.1016/j.cell.2016.08.072

[66] H. Li, X.X. Wang, B. Wang, L. Fu, G. Liu, Y. Lu, M. Cao, H. Huang, B. Javid, Latently and uninfected healthcare workers exposed to TB make protective antibodies against Mycobacterium tuberculosis, Proc. Natl. Acad. Sci. U. S. A., 114 (2017) 5023–5028. 10.1073/pnas.1611776114

[67] E.B. Irvine, A. O’Neil, P.A. Darrah, S. Shin, A. Choudhary, W. Li, W. Honnen, S. Mehra, D. Kaushal, H.P. Gideon, J.L. Flynn, M. Roederer, R.A. Seder, A. Pinter, S. Fortune, G. Alter, Robust IgM responses following intravenous vaccination with Bacille Calmette-Guerin associate with prevention of Mycobacterium tuberculosis infection in macaques, Nat. Immunol., (2021). 10.1038/s41590-021-01066-1

[68] M.F. Bachmann, G.T. Jennings, Vaccine delivery: a matter of size, geometry, kinetics and molecular patterns, Nature reviews. Immunology, 10 (2010) 787–796. 10.1038/nri2868

[69] T.N. Ellis, M.J. Kuehn, Virulence and immunomodulatory roles of bacterial outer membrane vesicles, Microbiology and molecular biology reviews : MMBR, 74 (2010) 81–94. 10.1128/MMBR.00031-09

