## Supplementary Information for "Immunomodulatory effect of mycobacterial outer membrane vesicles coated nanoparticles"

[&]Equal contribution

[#] Present Affiliation: Aten Porus Lifesciences Pvt Ltd, Bangalore, India

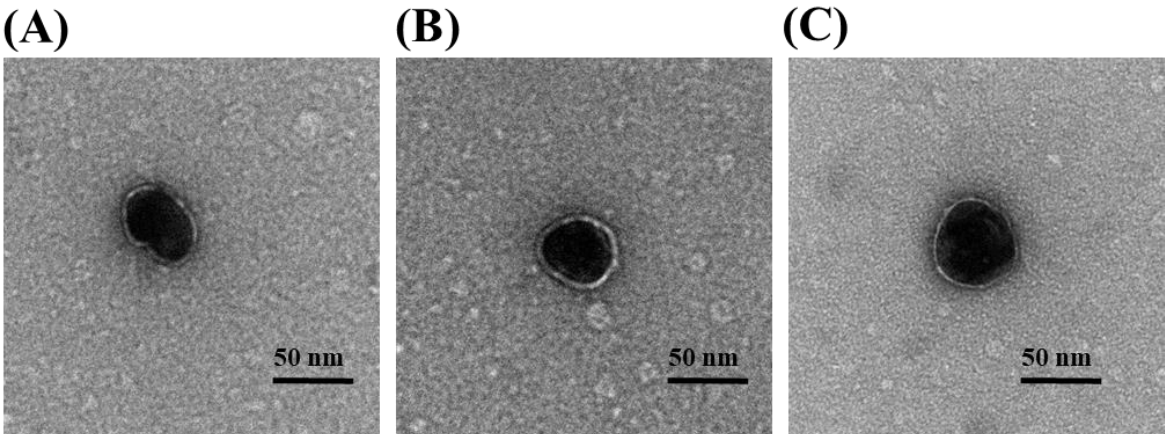


**Supplementary Figure 1:** TEM images of OMV-AuNP. Scale bar: 50 nm.

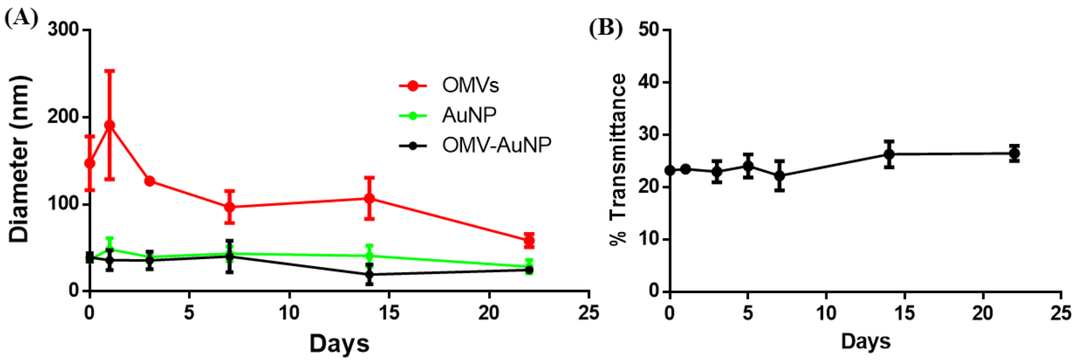


**Supplementary Figure 2: (A)** Size stability of gold nanoparticles (AuNP), free OMVs, or OMV-AuNP nanoparticles measured using DLS. **(B)** Transmittance values of OMV-AuNP solution measured by absorbance at 560 nm. Data in the graph represent the mean ± s.d.

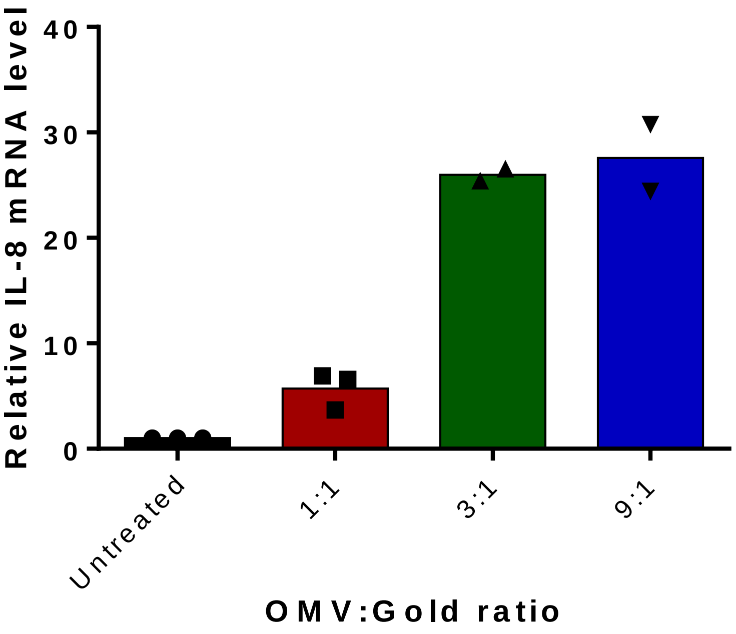


**Supplementary Figure 3:** Level of IL-8 mRNA when THP-1 cells were treated with OMV-AuNPs formed after extruding different ratios of OMV and gold nanoparticles. All conditions had 20 µg/mL gold nanoparticles extruded with different concentration of proteins present on OMVs.

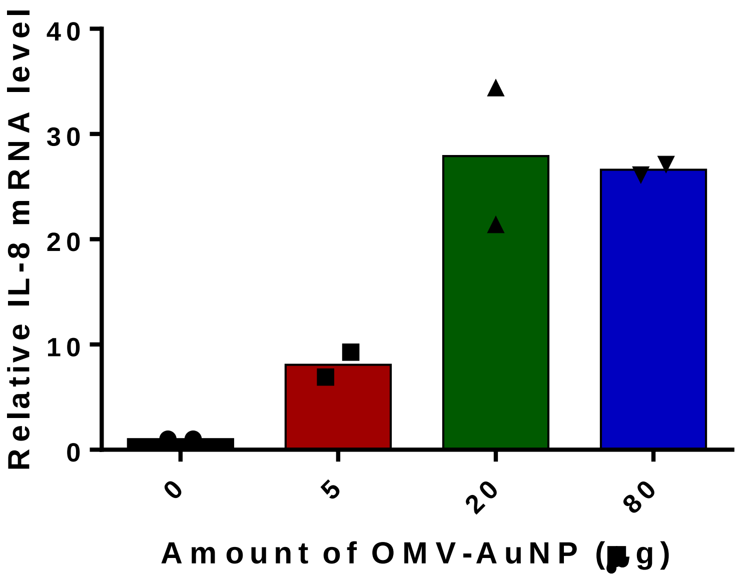


**Supplementary Figure 4:** Level of IL-8 mRNA when THP-1 cells were treated with increasing concentration of gold nanoparticles coated with OMVs.

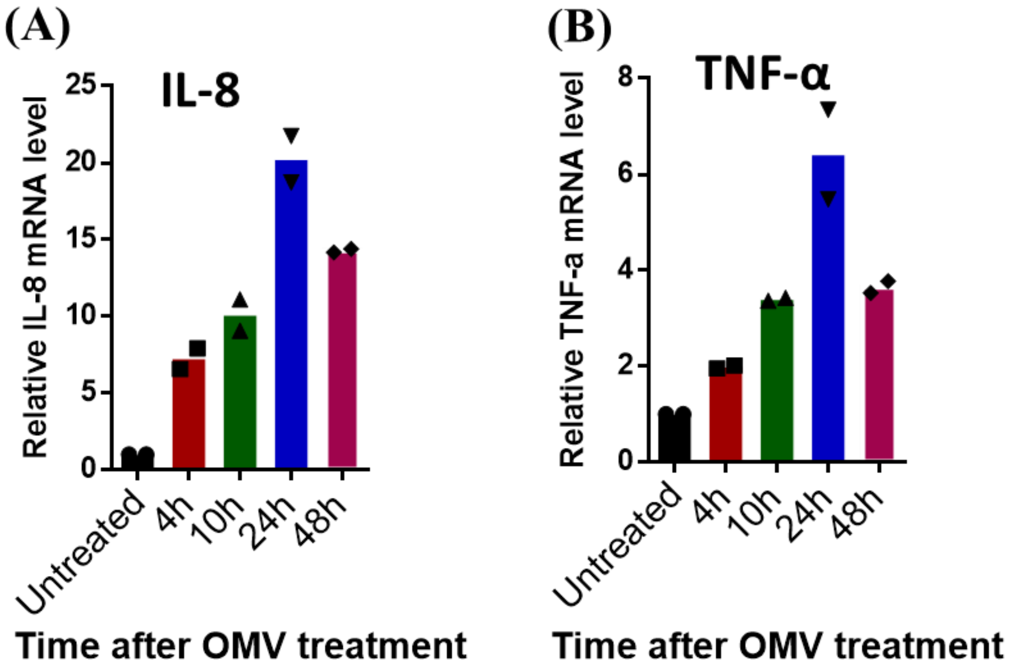


**Supplementary Figure 5:** THP1 cells treated with OMV-AuNP showed upregulation of (**A**) hIL-8 and (**B**) hTNF-α mRNA levels at various time points.

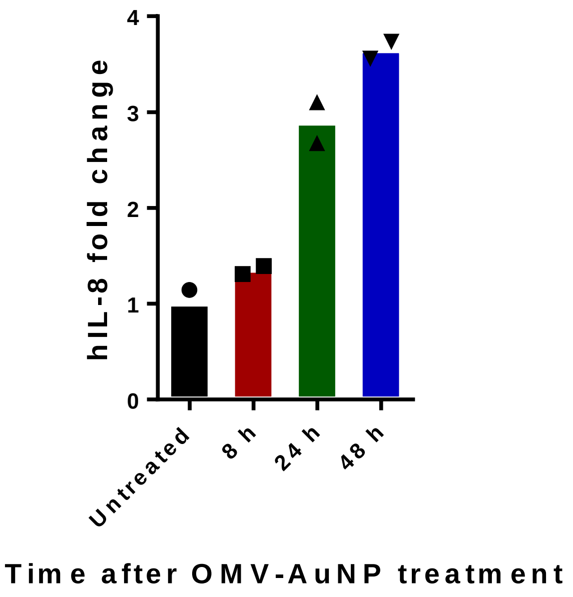


**Supplementary Figure 6:** The fold change in concentration of human IL-8 cytokine released into the media by THP-1 cells after treatment with OMV-AuNP compared to untreated cells at various time points as determined by ELISA.

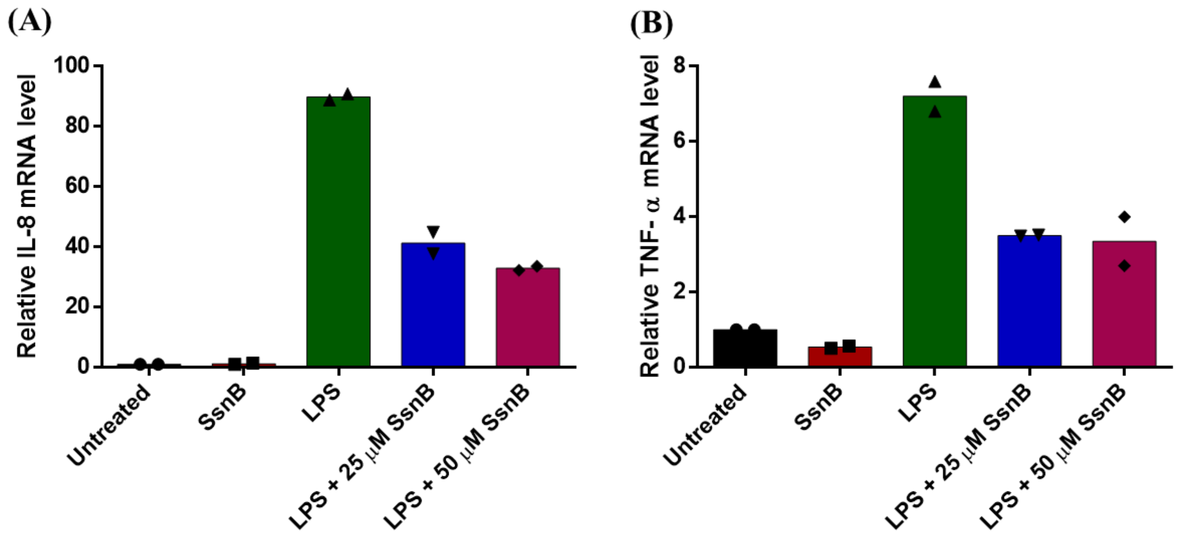


**Supplementary Figure 7:** mRNA level for cytokines when optimizing the Sparstolonin B (SsnB) concentration. THP-1 cells treated with varying concentrations of SsnB showed pronounced mRNA levels for cytokines (**A**) hIL-8 and (**B**) hTNF-α in the absence of SsnB when compared to cells treated with varying concentrations of SsnB.

**Supplementary Table 1: List of primers used for qPCR assay**

| Gene name | Forward Primer | Reverse Primer |
|---|---|---|
| hTNF-α | TCTTCTCGAACCCCGAGTGA | TATCTCTCAGCTCCACGCCA |
| hIL-8 | GAGAGTGATTGAGAGTGGACCAC | CACAACCCTCTGCACCCAGTTT |
| mTNF-α | GGTGCCTATGTCTCAGCCTCTT | GCCATAGAACTGATGAGAGGGAG |
| mIL-6 | TACCACTTCACAAGTCGGAGGC | CTGCAAGTGCATCATCGTTGTTC |
| mIL-1β | TGGACCTTCCAGGATGAGGACA | GTTCATCTCGGAGCCTGTAGTG |

**Supplementary Table 2: Total proteins identified from free OMVs and OMV-AuNS samples after LC-MS analysis**

| Protein IDs | H37Rv ortholog | Functional category | Function |
|---|---|---|---|
| MSMEG_6195 | Rv3680 | cell wall and cell processes | Probable anion transporter ATPase |
| MSMEG_6120 | #N/A | cell wall and cell processes | ABC transporter |
| MSMEG_3637 | Rv1842c | cell wall and cell processes | Conserved hypothetical membrane protein |
| MSMEG_3618 | Rv1860 | cell wall and cell processes | fibronectin attachment protein immunogenic protein MPT32 |
| MSMEG_6802 | #N/A | cell wall and cell processes | ABC transporter |
| MSMEG_6269 | Rv0362 | cell wall and cell processes | Possible Mg2+ transport transmembrane protein MgtE |
| MSMEG_1382 | #N/A | cell wall and cell processes | MmpL5 protein |
| MSMEG_3374 | #N/A | conserved hypotheticals | Unknown |
| MSMEG_2226 | #N/A | conserved hypotheticals | Unknown |
| MSMEG_6675 | #N/A | conserved hypotheticals | Unknown |
| MSMEG_1554 | #N/A | conserved hypotheticals | ethanolamine ammonia-lyase |
| MSMEG_5582 | #N/A | conserved hypotheticals | Unknown |
| MSMEG_1674 | #N/A | conserved hypotheticals | Unknown |
| MSMEG_0601 | #N/A | conserved hypotheticals | unknown |
| MSMEG_4574 | Rv2415c | conserved hypotheticals | Conserved hypothetical protein |
| MSMEG_5472 | Rv0992c | conserved hypotheticals | Conserved hypothetical protein |
| MSMEG_6551 | #N/A | conserved hypotheticals | hypothetical protein |
| MSMEG_1365 | Rv0652 | information pathways | 50S ribosomal protein L7/L12 RplL (SA1) |

| | | | |
|---|---|---|---|
| MSMEG_1368 | Rv0668 | information pathways | DNA-directed RNA polymerase (beta' chain) RpoC (transcriptase beta' chain) (RNA polymerase beta' subunit). |
| MSMEG_6927 | Rv3908 | information pathways | Possible mutator protein MutT4 |
| MSMEG_1721 | Rv1764, Rv3187, Rv3475 | Insertion seqs and phages | IS1137, transposase orfB |
| MSMEG_2681 | Rv0840c | intermediary metabolism and respiration | Probable proline iminopeptidase Pip (prolyl aminopeptidase) (pap) |
| MSMEG_0925 | #N/A | intermediary metabolism and respiration | SAM-dependent methyltransferase |
| MSMEG_3306 | #N/A | intermediary metabolism and respiration | zinc-binding alcohol dehydrogenase |
| MSMEG_4862 | #N/A | intermediary metabolism and respiration | 3-alpha-hydroxysteroid dehydrogenase |
| MSMEG_0306 | Rv3566c | intermediary metabolism and respiration | Arylamine N-acetyltransferase Nat (arylamine acetylase) |
| MSMEG_0485 | #N/A | intermediary metabolism and respiration | Amidase |
| MSMEG_0988 | Rv0534c | intermediary metabolism and respiration | 1,4-dihydroxy-2-naphthoate octaprenyltransferase MenA (DHNA-octaprenyltransferase) |
| MSMEG_0497 | #N/A | intermediary metabolism and respiration | glycerol dehydratase large subunit |
| MSMEG_1112 | #N/A | intermediary metabolism and respiration | aconitate hydratase, putative |
| MSMEG_6646 | #N/A | intermediary metabolism and respiration | methylisocitrate lyase |
| MSMEG_1702 | Rv3306c | intermediary metabolism and respiration | Probable amidohydrolase AmiB1 (aminohydrolase) |
| MSMEG_6030 | #N/A | intermediary metabolism and respiration | cytochrome p450 |
| MSMEG_2462 | #N/A | intermediary metabolism and respiration | carbon-monoxide dehydrogenase |

| | | | |
|---|---|---|---|
| MSMEG_3139 | Rv1472 | lipid metabolism | Possible enoyl-CoA hydratase EchA12 (enoyl hydrase) |
| MSMEG_1140 | Rv0564c | lipid metabolism | Probable glycerol-3-phosphate de-hydrogenase |
| MSMEG_0372 | Rv0242c | lipid metabolism | Probable 3-oxoacyl-[acyl-carrier protein] reductase FabG4 (3-ke-toacyl-acyl carrier protein reduc-tase) |
| MSMEG_2489 | #N/A | regulatory proteins | transcriptional regulator, GntR |
| MSMEG_0296 | #N/A | regulatory proteins | transcriptional regulator, MarR family protein |
| MSMEG_1369 | #N/A | regulatory proteins | LacI-family protein transcriptional regulator |
| MSMEG_6376 | #N/A | regulatory proteins | transcriptional regulator LacI fam-ily protein |
| MSMEG_1582 | Rv3418c | virulence, detoxification, ad-aptation | 10 kDa chaperonin GroES (protein CPN10) (protein GroES) (BCG-a heat shock protein) (10 kDa anti-gen) |
| MSMEG_6821 | #N/A | virulence, detoxification, ad-aptation | NLP/P60 family protein (pepti-doglycan hydrolase) |
